# Inhibition of V-ATPase function drives apoptosis via GCN1/GCN2 kinase signaling

**DOI:** 10.64898/2026.03.27.714872

**Authors:** Filip Gallob, Severin Lechner, Dominika Tučková, Petra Divoká, Yevheniia Tyshchenko, Danica Drpić, Jan Hájek, Lukas Englmaier, Kateřina Delawská, Miriam Unterlass, Mariana Araujo, Georg E. Winter, Pavel Hrouzek, Andreas Villunger

**Author notes:** **Correspondence:** Prof. Andreas Villunger, PhD, Institute for Developmental Immunology Biocenter, Medical University of Innsbruck Innrain 80, A-C020, Innsbruck, AT, Ph: +43-512-S003-70380 Fax: +43-512-S003-73SC0.

## Abstract

Natural products are a rich source of bioactive molecules that have served both as templates for drug discovery and as tools to uncover fundamental biological processes. While characterizing the pro-apoptotic activity of the cyanobacterial metabolite Nostatin A, we identified vacuolar-type H⁺-ATPase (V-ATPase) as its molecular target and uncovered an unexpected signalling response preceding cell death initiation. V-ATPase inhibition rapidly activates the integrated stress response (ISR) through engagement of the GCN1/GCN2 kinase module, indicative of ribosomal collisions and translational shutdown. This response is conserved across established V-ATPase inhibitors, including bafilomycin A1, but not with compounds disrupting lysosomal function by other means. Mechanistically, V-ATPase inhibition depletes the pro-survival protein MCL-1 resulting in BAX/BAK-dependent mitochondrial apoptosis. Loss of MCL-1 creates a vulnerability that renders cells dependent on co-expressed BCL-2 family proteins, enabling potent synergy with the BH3 mimetics ABT-737 or venetoclax. Taken together, our results reveal a therapeutically exploitable vulnerability in V-ATPase–reliant or MCL-1 dependent cancers.

## Introduction

Natural products (NPs) and their structural derivatives play a central role in shaping modern therapeutics and guide drug discovery. Many compounds derived from natural sources are now used as standard-of-care therapeutics across diverse clinical indications, including statins for cardiovascular disease, immunosuppressive agents such as rapamycin for organ transplantation, and taxanes for cancer treatment^1,2^. Beyond their direct therapeutic applications, natural products serve as powerful molecular probes to discover new biology^3^.

A number of landmark discoveries in cell biology were enabled through the study of natural products. Characterization of the molecular targets of rapamycin, cyclosporin A, and FK506 (tacrolimus), for example, not only transformed immunosuppressive therapy but, also revealed critical signaling pathways beyond the immune system. Rapamycin led to the discovery of the mTOR pathway, a central regulator of cell growth and metabolism, while cyclosporin A and FK506 uncovered key aspects of calcium-dependent signaling through inhibition of the phosphatase calcineurin^4,5^. These examples illustrate how natural products have not only expanded the therapeutic arsenal but also provided key insights into cell biology, often functioning as the first described molecular glues^6^.

Among the expanding families of natural products are ribosomally synthesized and post-translationally modified peptides (RiPPs), which display a remarkable diversity of biological activities^7^. RiPPs are produced by bacteria and fungi as precursor peptides that undergo extensive post-translational tailoring to yield highly structured bioactive molecules. These modifications commonly include macrocyclization and the formation of thiazole and oxazole heterocycles. Together, these structural features enhance conformational stability, improve target recognition, and increase resistance to proteolytic degradation^8^.

Peptide-based natural products such as RiPPs are attractive scaffolds for drug development^9^. In contrast to many small molecules, peptide frameworks are readily amenable to rational chemical modification, enabling optimization of pharmacokinetic properties, stability, and target selectivity^9^. Advances in synthetic chemistry and peptide engineering have further expanded the possibilities for modifying these scaffolds, facilitating the development of peptide-based therapeutics with improved efficacy and reduced toxicity^10,11^.

Among widely studied drug targets that still lack effective inhibitors for clinical use is the vacuolar-type H⁺-ATPase (V-ATPase)^12^. V-ATPase is a multi-subunit protein complex responsible for acidifying intracellular compartments such as lysosomes, endosomes, the Golgi apparatus, and secretory vesicles^13^. By pumping protons (H⁺) across membranes, V-ATPase generates electrochemical gradients that support numerous cellular processes. One of its primary functions is the acidification of lysosomes, which is required for the degradation of macromolecules during lysosomal protein turnover and autophagy-mediated cargo recycling. Through this activity, lysosomal V-ATPase plays a central role in maintaining cellular homeostasis by removing damaged organelles, aggregated or redundant cellular components^14^. In endosomes, acidification supports the sorting and trafficking of internalized molecules, thereby ensuring proper cellular signaling and resource allocation^15^. Beyond these roles, V-ATPase also influences cellular growth, metabolism, and differentiation through its contribution to cytosolic pH regulation, affecting processes such as protein trafficking, neurotransmitter release, and insulin secretion^16^.

Given its central role in lysosomal function and metabolic adaptation, V-ATPase has emerged as an attractive therapeutic target. Dysregulation of V-ATPase activity has been implicated in several human diseases, renal tubular acidosis, cancer, and neurodegenerative disorders^13,17,18^. In cancer cells in particular, lysosomal pathways and autophagy often play critical roles in sustaining growth and survival under metabolic stress. Inhibition of V-ATPase disrupts lysosomal degradation and autophagic flux, leading to accumulation of damaged proteins and organelles, metabolic stress, and ultimately cell death. Despite this strong therapeutic rationale, the clinical development of V-ATPase inhibitors has been limited by systemic toxicity^12^.

Although V-ATPase inhibition is widely known to induce cell death, the molecular pathways linking impaired lysosomal acidification to activation of the apoptotic machinery remain ill-defined^13^. Several mechanisms have been proposed, including accumulation of autophagosomes and undegraded cargo, induction of endoplasmic reticulum (ER) stress, increased reactive oxygen production, and mitochondrial dysfunction^19^. These events are thought to converge on mitochondrial outer membrane permeabilization, regulated by the BCL-2 family of proteins, ultimately leading to caspase activation^12^. In this framework, apoptosis following V-ATPase inhibition has largely been interpreted as a secondary consequence of disrupted autophagy and unfolded protein stress signaling, but not as a direct consequence of V-ATPase impairment.

A known consequence of lysosomal perturbation is the reduced replenishment of cytosolic free amino acids derived from lysosomal degradation, which can limit protein synthesis^20^. However, how cells sense and respond to such disturbances in nutrient supply, remains incompletely understood. One pathway linking metabolic stress to cell fate decisions is the integrated stress response (ISR), centered around the transcription factor ATF4^21^. ISR activation occurs through phosphorylation of the translation initiation factor eIF2α on serine 51 by the kinases GCN2, HRI, PERK, or PKR, which suppresses global protein synthesis while selectively enhancing ATF4 translation. This leads to the induction of a transcriptional program that promotes metabolic adaptation by regulating amino acid metabolism, redox balance, and stress-response pathways^22^. While ATF4 initially supports adaptive responses that restore cellular homeostasis, prolonged ISR signaling can shift this program toward apoptosis through induction of pro-apoptotic BCL-2 family members^23^. A less well understood consequence of ISR activation is the loss of the short-lived pro-survival BCL-2 family member MCL-1 and the extent to which this contributes to mitochondrial apoptosis^24,25^. Whether V-ATPase inhibition and, by extension, lysosomal dysfunction engage this pro-apoptotic signaling axis remains unresolved.

In this study, we investigated the mode of action (MOA) of the previously uncharacterized cyanobacterial RiPP Nostatin A^26^. Through target-agnostic analysis of NoA-induced cytotoxicity, transcriptomic analyses and subsequent chemoproteomic target identification, we uncover an unexpected link between V-ATPase inhibition, activation of the ISR, and mitochondrial apoptosis. We show that NoA gains access to lysosomal V-ATPase through endolysosomal pathways and inhibits lysosomal acidification. This in turn leads to rapid GCN1/GCN2-dependent activation of the ISR and apoptosis. Cell death following V-ATPase inhibition is largely driven by the loss of pro-survival MCL-1, rather than BH3 only protein activation. Importantly, this mechanism can be generalized to other dedicated V-ATPase inhibitors, and we demonstrate synergy between inhibition of V-ATPase and pharmacological targeting of BCL-2 family proteins in tumor cell killing. Together, these findings not only reveal an unexpected pro-apoptotic signaling cascade downstream of V-ATPase inhibition but also highlight Nostatin A as a promising lead compound that warrants further investigation as a potential cancer therapeutic.

## Results

### Nostatin A induces mitochondrial apoptosis across different cancer cell lines

NoA has been assigned potent cytostatic activity in the low nanomolar range in a panel of cancer cell lines^26^(supp. Fig. 1a). Here, we continued to characterize its impact on cell growth and survival using three different human model cell lines: myeloid leukemia-derived near-haploid HAP-1 cells, pre-B acute lymphoblastic leukemia Nalm-6 cells and colorectal cancer cells (CRC) HCT-116. While HAP-1 cells are deficient for the tumor suppressor p53, Nalm-6 and HCT-116 cells are p53 proficient.

We started with characterizing the cell death kinetics and metabolic activity upon NoA treatment by using the cell titer glow (CTG) assay, which assesses cellular ATP levels as a proxy for viability. Exposing HAP-1, Nalm-6 and HCT-116 cells to graded doses of NoA promoted a dose-dependent decrease in ATP content after 72h (Fig. 1a, left panel, supp. Fig. 1b & c). We previously found, that loss of the proapoptotic BCL-2 family proteins BAX/BAK (BB dKO), or inhibition of caspases with the pan-caspase inhibitor Q-VD-OPH caused an increase in IC50 values in our initial report on NoA-induced cytotoxicity in HeLa cells^26^. To confirm these findings across our panel of cell lines, we employed derivatives deficient in BAX and BAK, dubbed BB dKO (Fig 1a, right panel)^27^. We observed that BAX and BAK co-deletion or chemical caspase inhibition led to a modest but significant increase of IC50 values in HAP-1 cells (Fig. 1a), similar to what we noted previously^26^. However, Nalm-6 and HCT-116 cells deficient for BAX and BAK did not show a significant change in IC50 values in response to NoA, suggesting a general metabolic shut down (supp. Fig 1b & c). We then characterized the kinetics of the loss in metabolic activity in HAP-1 cells at 0.5, 1 and 2x the IC50 of NoA. Treating HAP-1 WT and BB dKO cells at these concentrations revealed that NoA promoted a time and dose-dependent decrease in ATP in both genotypes, albeit less pronounced in the absence of BAX and BAK (Fig. 1b), indicative of prolonged cell survival.

**Figure 1:**
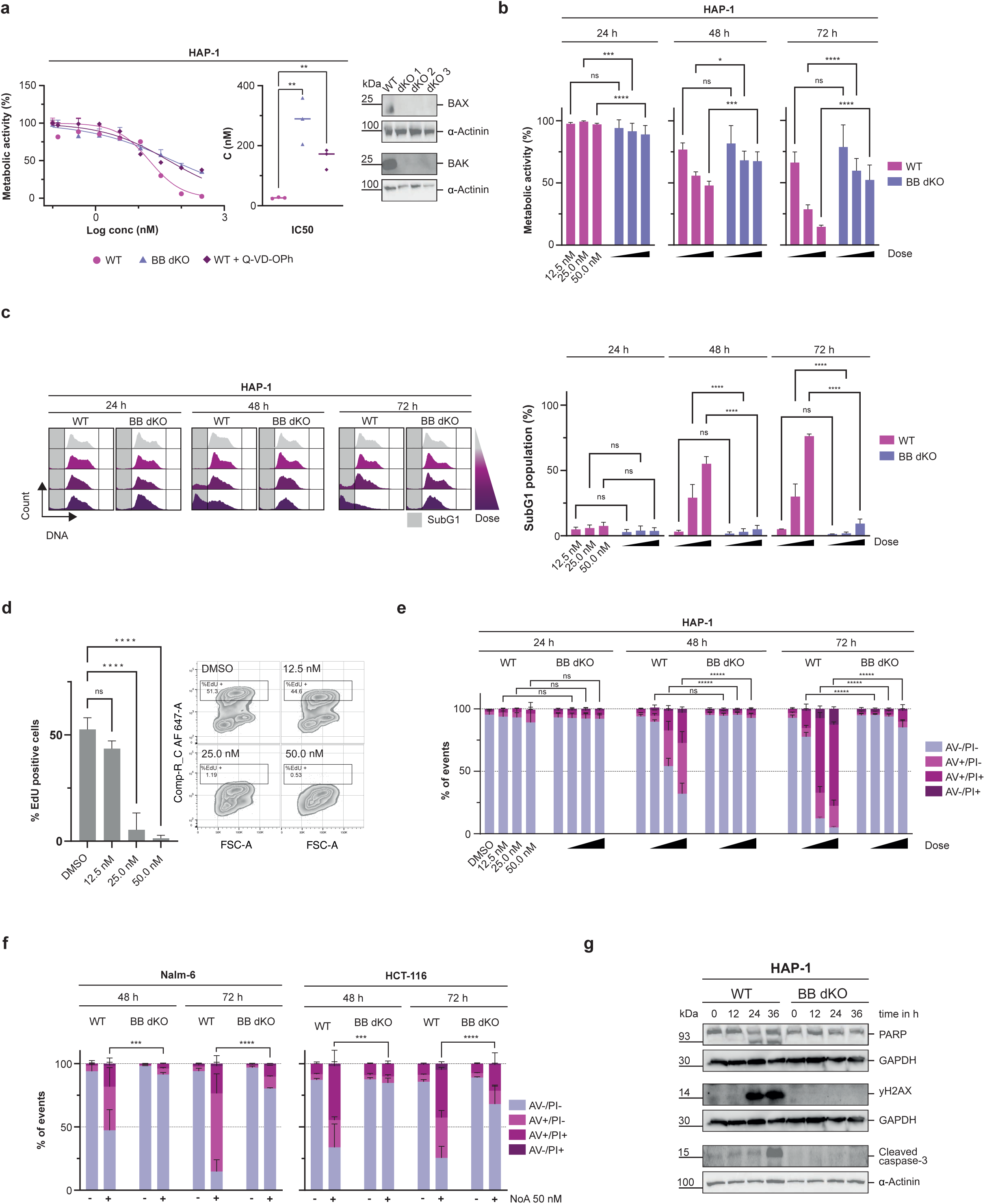
Nostatin A induces mitochondrial apoptosis across different cancer cell lines. a. Left: Metabolic activity of HAP-1 WT and HAP-1 BAX/BAK dKO cells treated with graded concentrations of NoA alone or in combination with the pan-caspase inhibitor Q-VD-OPh (10 μM) for 72 h was assessed using the CellTiter-Glo® Luminescent Cell Viability Assay. Data points represent mean ± SD of three technical replicates. IC₅₀ values are shown as mean ± SD of N = 3 biological replicates and were determined by linear regression. Statistical significance between IC₅₀ values was assessed using an unpaired t-test. \*\**P* < 0.05; Right: Validation of BAX- and BAK-deficient HAP-1 cells by western blot. b. Metabolic activity of HAP-1 WT and HAP-1 BB dKO cells treated with 12.5, 25, or 50 nM NoA was measured after 24, 48, and 72 h using the CellTiter-Glo® Luminescent Cell Viability Assay. Bars represent mean ± SD of three biological replicates, each with three technical replicates. Metabolic activity was normalized to DMSO controls at each time point. Statistical significance was determined using two-way ANOVA with Bonferroni correction for multiple comparisons, comparing WT and BB dKO cells at corresponding concentrations and time points. \**P* < 0.05, \*\*\**P* < 0.001, \*\*\*\**P* < 0.0001 c. Left: Representative cell cycle profiles of HAP-1 WT and BB dKO cells after exposure to 12.5, 25 and 50 nM of NoA. Right: Quantification of the SubG1 population represented as bar graphs, showing means and SD of N=3 biological replicates. Statistical significance was determined using a two-way ANOVA with Bonferroni correction for multiple comparisons, comparing SubG1 percentages of corresponding concentrations of WT with BB dKO cells at every time point. \*\*\*\**P* < 0.0001 d. EdU incorporation in HAP-1 WT celle after 24 h treatment with graded concentrations of NoA. Bar graphs (left) are showing means ± SD of N=3 biological replicates of % of EdU positive cells with representative dot plots (right). Statistical significance was assessed using a one-way ANOVA with Dunnet’s multiple comparisons test. \*\*\*\**P* < 0.0001 e. HAP-1 WT and HAP-1 BB dKO cells were treated with either DMSO or 12.5, 25 and 50 nM of NoA. Annexin V surface binding and Propidium Iodide uptake was measured after 24, 48 and 72 h to assess viability of the cells. Bar plots show mean ± SD of three biological replicates. Legend: AV−/PI− = live cells, AV+/PI− = early apoptosis, AV+/PI+ and AV−/PI+ = late apoptosis. Statistical significance was determined using a two-way ANOVA with Bonferroni correction for multiple comparisons, comparing percentages of live cells of corresponding concentrations between WT and BB dKO cells at every time point. \*\*\*\**P* < 0.0001 f. HCT-116 and Nalm-6 WT and BB dKO cells were treated with either DMSO or 50nM of NoA. Annexin V surface binding and Propidium Iodide uptake was measured after 48 and 72 h to assess viability of the cells. Bar plots show mean ± SD of three biological replicates. Legend: AV−/PI− = live cells, AV+/PI− = early apoptosis, AV+/PI+ and AV−/PI+ = late apoptosis. Statistical significance was determined using a two-way ANOVA with Bonferroni correction for multiple comparisons, comparing percentages of live cells of corresponding concentrations between WT and BB dKO cells at every time point. \*\*\**P* < 0.0005, \*\*\*\**P* < 0.000. g. Western blot analysis of HAP-1 WT and HAP-1 BB dKO cells treated with 50 nM NoA. Cells were harvested at 0, 12, 24, and 36 h after treatment, and protein lysates were probed with the indicated antibodies.

Cell cycle profiling by flow cytometric DNA content analysis revealed that cells responded initially with a G1 arrest to NoA treatment, before engaging a cell death response starting after 24h of treatment as evidenced by an increase in cells displaying reduced DNA content (SubG1 fraction) (Fig. 1c). While HAP-1 WT cells treated with 1 or 2x IC50 concentrations show a similar impact on their cell cycle profiles, HAP-1 cells treated at 0.5 IC50 were able to maintain a normal cell cycle profile. Although HAP-1 BB dKO cells did not show an increase in the SubG1 population, indicating protection from cell death, they also failed to re-enter the cell cycle after prolonged NoA treatment and remained arrested in G1 phase at higher doses (Fig. 1c). We additionally performed an EdU incorporation assay in HAP-1 WT cells to confirm that increasing NoA concentrations lead to gradual decrease in S-phase activity, underscoring that a G1 arrest after NoA treatment precedes apoptosis induction (Fig. 1d).

To establish the timepoint when cells become apoptotic, Annexin V surface binding was assessed over time at different doses of NoA. Consistent with the cell cycle profiling data, we did not observe significant Annexin V cell surface binding at 24h, suggesting induction of cell death between 24 and 48h of treatment (Fig. 1e). Annexin V binding was completely abolished in the BAX/BAK deficient HAP-1 cells at all time points, in line with their solid G1 arrest and a lack of cells accumulating in the SubG1 gate (Fig. 1c). We also confirmed BAX- and BAK-dependent induction of apoptosis following NoA treatment in Nalm-6 and HCT-116 cells, as BB dKO cells showed strongly reduced Annexin V surface binding (Fig. 1f). Finally, western blot analysis confirmed that caspase-activation and apoptosis-associated DNA damage, respectively, as shown by monitoring for PARP1 cleavage or γH2AX phosphorylation, respectively, are downstream of BAX and BAK activation (Fig. 1g). Similar findings were made in Nalm-6 and HCT-116 cells (supp. Fig. 1d). Furthermore, both cell lines responded with an initial G1 arrest to NoA treatment before an increase in the SubG1 population was observed (supp. Fig. 1e). BAX- and BAK-deficient clones, however, showed a significantly reduced SubG1 population compared to WT cells (supp. Fig 1f). Exploiting a series of gene-edited Nalm-6 cells, lacking different caspase family members, confirmed the induction of mitochondrial apoptosis as the main cell death response engaged in response to NoA treatment. Loss of Caspase-9 or combined loss of effector Caspases-3/6/7 were providing a similar degree of cell death protection as did loss of BAX and BAK, while loss of initiator caspase-2 or -8 had no effect (supp. Fig 1e and f).

### Nostatin A induces ATF4- and JUN-dependent transcriptome changes preceding apoptosis

We next sought to identify the pro-apoptotic signals responsible for the observed cell death phenotype. As no increase in Annexin V surface binding or SubG1 population was detected at 12 h or 24 h after treatment, we concluded that most cells had not yet activated caspases or committed to apoptosis at these time points. To identify signaling pathways that may drive intrinsic apoptosis within this window, we performed transcriptomic analysis of HAP-1 cells.

A GO term enrichment analysis of differentially expressed genes (DEGs) revealed that ER stress response genes were the most significantly upregulated after 12 h of NoA exposure (Fig. 2a), followed by genes involved in cholesterol/sterol synthesis and ER-to-Golgi transport. The most strongly downregulated terms at this time point were associated with ncRNA processing and ribosome biogenesis. Notably, downregulation of these pathways became substantially more pronounced at 24 h, suggesting a translational shutdown preceding cell death initiation and indicative of proteotoxic stress (supp. Fig. 2a).

**Figure 2:**
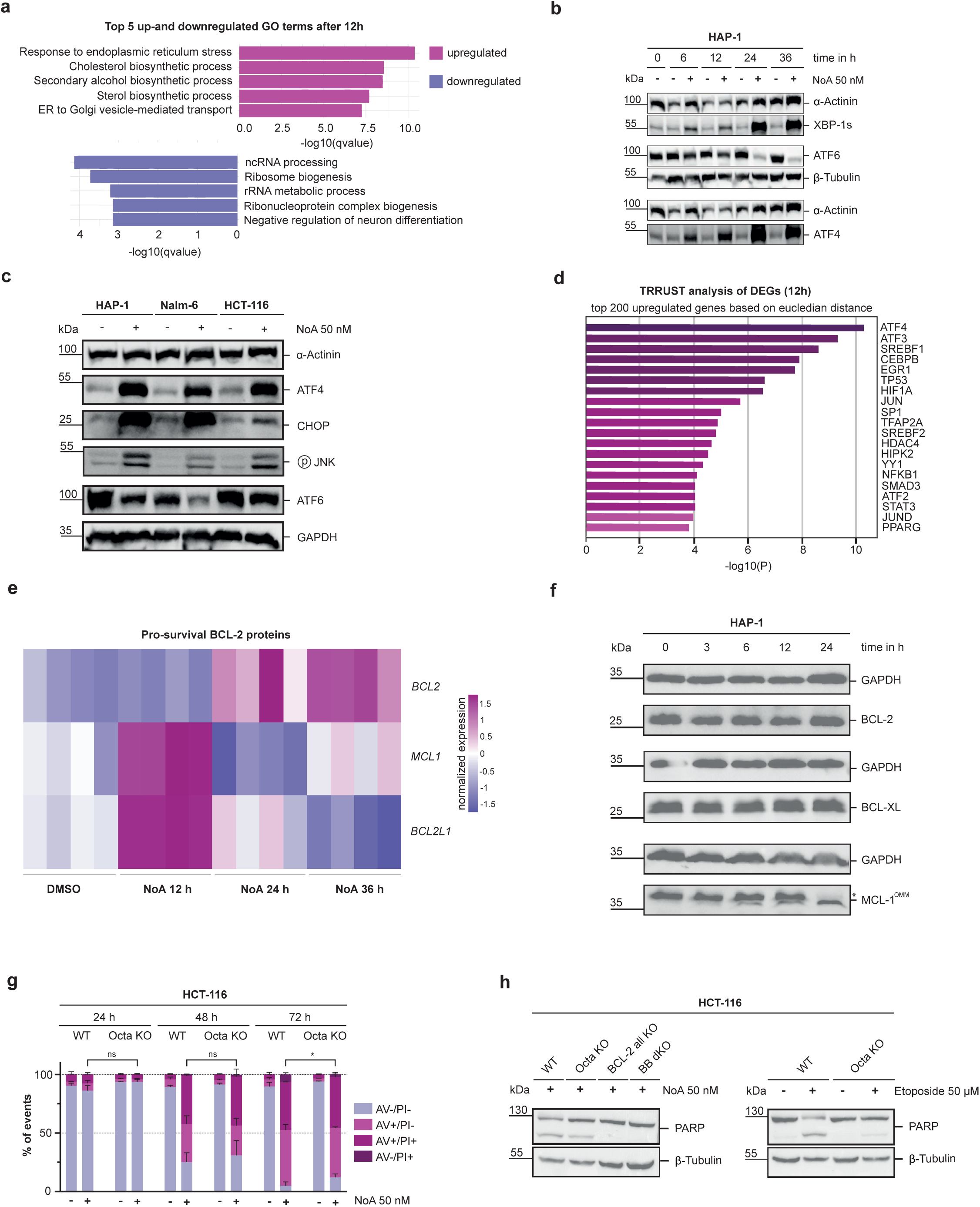
Nostatin A induces ATF4- and JUN-dependent transcriptome changes preceding apoptosis. a. Gene ontology analysis of differentially expressed genes of HAP-1 cells after 12 h treatment with 50 nM of NoA or DMSO vehicle control. b. Western blot analysis of HAP-1 WT cells treated with DMSO or 50 nM NoA. Cells were harvested at 0, 12, 24, and 36 h after treatment, and protein lysates were probed with the indicated antibodies. c. Western blot analysis showing activation of canonical UPR signaling proteins in HAP-1, Nalm-6 and HCT-116 WT cells exposed to 50 nM of NoA or DMSO for 24 h. d. TRRUST analysis of differentially expressed genes in HAP-1 cells treated with 50 nM NoA for 12 h, showing transcription factors predicted to be responsible for the upregulation of the top 200 genes after treatment. e. Heatmap showing regulation of pro-apoptotic BCL-2 genes derived from the RNA-seq dataset. Values represent row-scaled, variance-stabilizing transformation (VST)–normalized expression of differentially expressed genes. Values of n=4 technical replicates are shown. f. Western blot analysis of HAP-1 WT cells treated with 50 nM NoA. Cells were harvested at 0, 3, 6, 12 and 24 h after treatment, and protein lysates were probed with the indicated antibodies. MCL-1 OMM refers to the full-length, anti-apoptotic isoform MCL-1. g. HCT-116 WT and Octa KO clones were treated with 50nM of NoA. Annexin V surface binding and Propidium Iodide uptake was measured after 24, 48 and 72 h to assess cell viability. Bar plots show mean ± SD of three biological replicates. Legend: AV−/PI− = live cells, AV+/PI− = early apoptosis, AV+/PI+ and AV−/PI+ = late apoptosis. Statistical significance was determined using a two-way ANOVA with Bonferroni correction for multiple comparisons, comparing percentages of live cells of corresponding concentrations between WT and Octa KO clones. \**P* < 0.05, h. Left: WB analysis of HCT-116 WT, Octa KO, BCL-2 all KO and BB dKO cells after 48 h of treatment with 50 nM of NoA. Right: Western blot analysis of HCT-116 WT and Octa KO after 48 h of treatment with 50 μM of Etoposide.

ER stress is canonically mediated by three signaling branches - ATF6, IRE1α, and PERK - collectively referred to as the unfolded protein response (UPR)^28^. To validate the transcriptomic findings and determine which of these pathways contributes to apoptosis induction, we performed western blot analysis in HAP-1 cells (Fig. 2b). We found that ATF4 and the active variant of XBP-1, XBP-1s, a downstream targets of both PERK and IRE1α signaling, showed pronounced stabilization, as early as 6h after treatment (Fig. 2b). Similar hallmarks of ER stress pathway activation were observed in Nalm-6 and HCT-116 cells, confirming robust induction of the UPR across NoA-treated cell lines, including an increased JNK phosphorylation, a known downstream consequence of IRE1α activation (Fig. 2c)^29^.

PERK and IRE1α indirectly regulate the transcription factors ATF4 and JUN through phosphorylation of eIF2α and JNK, respectively^21,28^. Sustained activation of these stress pathways induces pro-apoptotic BCL-2 family proteins, including the BH3-only proteins *BBC3*/PUMA and *PMAIP1*/NOXA downstream of PERK–ATF4 signaling, and *BCL2L11*/BIM downstream of IRE1α–JNK signaling (supp. Fig. 2b)^30^. To confirm that ATF4- and JUN-regulated genes are induced after 12 h of NoA treatment, we performed TRRUST analysis (Transcriptional Regulatory Relationships Unraveled by Sentence-based Text Mining) to identify transcription factors driving gene expression changes leading to mitochondrial apoptosis. This analysis identified ATF4, and to a lesser extent JUN, as key regulators (Fig. 2d).

We then mined the transcriptomics data set to confirm that canonical ATF4 and JUN target genes are indeed upregulated (supp. Fig. 2c and d)^31,32^. In addition, we confirmed that pro-apoptotic BH3-only proteins previously associated with the apoptotic arm of the UPR were upregulated on mRNA level. Indeed, *PUMA* and *NOXA* transcripts were found significantly increased after 24 h, while *BIM* was not (supp. Fig. 2e). To validate this on protein level, we performed western blot analysis and probed for BIM, PUMA and NOXA in HAP-1 cells. Consistent with our transcriptomics analysis we saw an upregulation of NOXA and PUMA on the protein level. We also observed an increase of proapoptotic BIM isoforms (supp. Fig. 2f).

Having established that BIM, NOXA, and PUMA are upregulated at the protein level following NoA treatment, we next quantified their relative contributions to NoA-induced apoptosis. To this end, Nalm-6 clones deficient in PUMA, NOXA, BIM, or both BIM and NOXA were generated, treated with NoA and cell survival was assessed by Annexin V/PI staining. While loss of NOXA or BIM alone did not confer protection, PUMA-deficient cells and cells lacking both BIM and NOXA displayed a modest survival advantage after 72 h of treatment (supp. Fig. 2g). However, this protection was markedly weaker than that observed upon loss of BAX and BAK (Fig. 1f), indicating substantial redundancy among BH3-only proteins, or alternative modes of BAX/BAK activation, such as the loss of prosurvival proteins. Along these lines, mining the RNA-seq data revealed dynamic regulation of pro-survival BCL-2 family members following NoA treatment. Notably, *MCL-1* transcripts showed an early transient increase followed by a marked decline at later time points (Fig. 2e). A similar pattern was observed for *BCL2L1/*BCL-XL, whereas *BCL-2* transcripts increased despite ongoing apoptosis induction. At the protein level, however, only the short-lived pro-survival factor MCL-1 showed a pronounced time-dependent decrease across HAP-1, HCT-116, and Nalm-6 cells, while BCL-XL and BCL-2 levels remained largely unchanged (Fig. 2f and supp. Fig. 2h)^33^. Notably, we observed a loss of the full-length anti-apoptotic isoform of MCL-1 localized to the mitochondrial outer membrane (dubbed MCL-1 OMM) in HAP-1 and HCT-116 cells, while we did not observe this N-terminally truncated isoform in Nalm-6 cells^34^.

To further dissect the contribution of pro-apoptotic BH3-only proteins versus loss of pro-survival BCL-2 family members, we employed HCT-116 cells lacking all eight pro-apoptotic BH3-only proteins (Octa KO) or cells deficient in all established BCL-2 family members (BCL-2 all KO)^35^. Previous work has shown that Octa KO cells remain susceptible to mitochondrial apoptosis following depletion of MCL-1 and BCL-XL through siRNAs^35^. Given the poor protection inferred by loss of BH3-only proteins, we wondered whether HCT-116 Octa KO cells remained sensitive to NoA when compared with BCL-2 all KO or BB dKO cells. Indeed, Octa KO cells displayed levels of cell death comparable to WT cells upon NoA treatment (Fig. 2g). Consistently, Octa KO cells also exhibited executioner caspase activation, as evidenced by PARP cleavage, which was absent in BCL-2 all KO and BAX/BAK dKO cells (Fig. 2h). Notably, Octa KO cells treated with the DNA-damaging agent etoposide showed minimal PARP cleavage, suggesting that NoA-induced apoptosis is largely driven by loss of pro-survival proteins rather than activation of BH3-only proteins (Fig. 2h).

### NoA activates the ISR and ribotoxic stress response through GCN1/GCN2 and ZAKα

Having established that NoA-treated cells exhibit hallmarks of ER stress and activate the UPR, we hypothesized that NoA promotes the accumulation of unfolded proteins. To test this, we measured protein unfolding using TPE-MI, a probe that becomes fluorescent upon binding of exposed cysteine residues in unfolded proteins^36^. Contrary to our expectation, NoA treatment did not increase the TPE-MI signal up to 12 h, whereas proteasome inhibition with MG132 or inhibition of protein glycosylation with tunicamycin robustly increased fluorescence (Fig. 3a). An increase in TPE-MI signal was only observed at later time points and at 2× IC50 NoA. Notably, transcriptomic analysis showed strong upregulation of ATF4 target genes already after 12 h of treatment, while WB analysis showed ATF4 upregulation as early as 6 h of treatment. Since unfolded proteins were not increased at this time point, we reasoned that UPR activation may represent a later event and that ATF4 induction might arise from a different trigger. In addition to PERK, the kinases GCN2, HRI, and PKR can phosphorylate eukaryotic initiation factor 2 alpha (eIF2α) and activate the integrated stress response (ISR). eIF2α phosphorylation suppresses global translation while promoting selective translation of stress response genes, prominently regulated by ATF4^21^.

**Figure 3:**
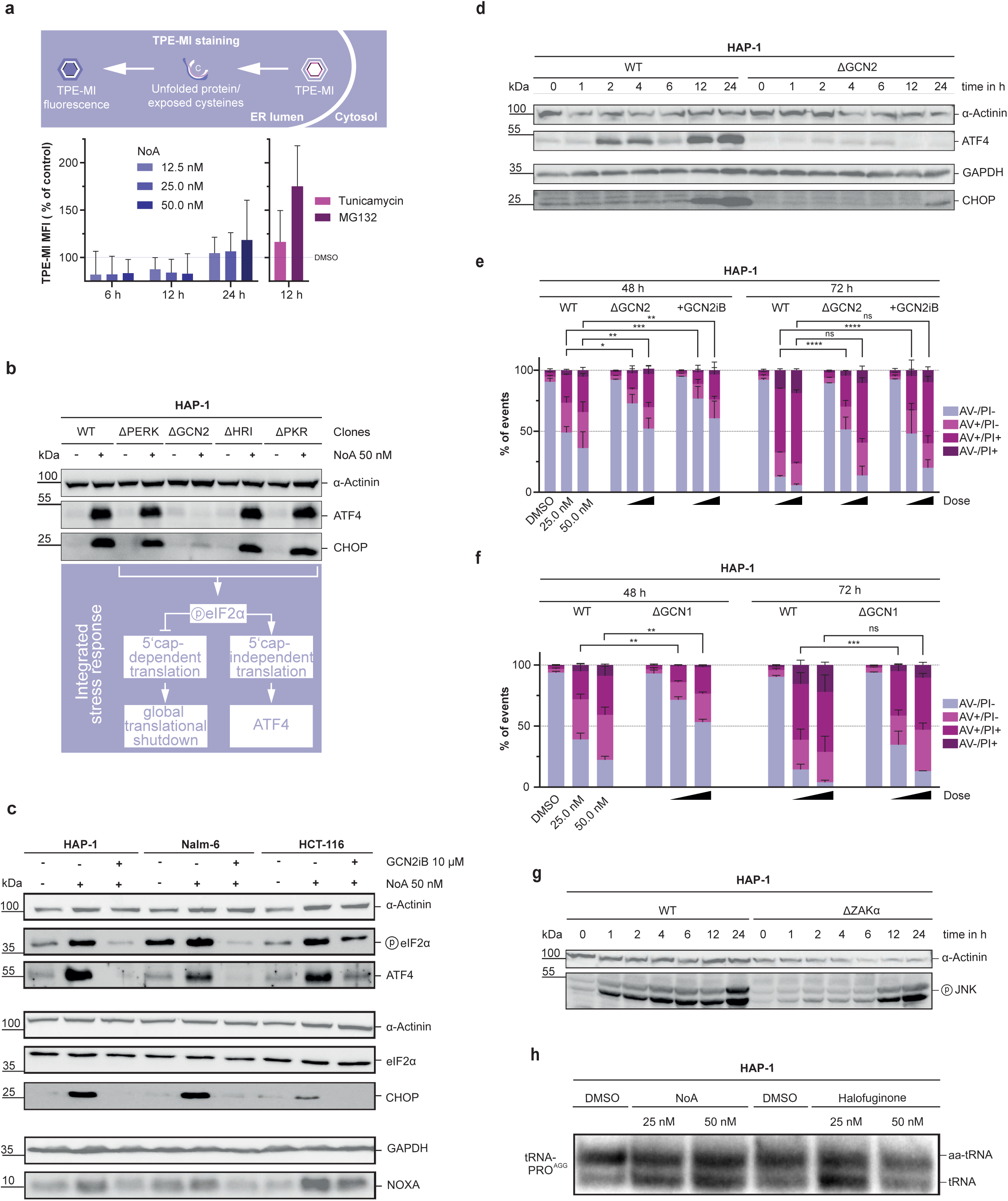
NoA activates the ISR and ribotoxic stress response through GCN1/GCN2 and ZAKα. a. TPE-MI fluorescence measured in HAP-1 WT cells using FACS analysis after treatment with graded concentrations of NoA or 500 nM of Tunicamycin or MG321. Bar plots show mean fluorescence intensity (MFI) of treated samples as percentage over DMSO treated control. Values show mean ± SD of three biological replicates. b. WB analysis of HAP-1 clones deficient for one of the four eIF2α kinases after a 24 h treatment with 50 nM of NoA. Scheme below illustrates the integrates stress response pathway. c. WB analysis comparing ISR activation in HAP-1, Nalm,6 and HCT-116 WT cells after treatment with 50 nM NoA alone or in combination with the GCN2 inhibitor GCN2iB (10 μM) after 24 hours. d. Time resolved WB analysis of HAP-1 WT and GCN2 KO cells after treatment with 50 nM of NoA. Lysates were probed with the indicated antibodies. e. HAP-1 WT and HAP-1 GCN2 KO cells were treated with 25 and 50 nM of NoA. HAP-1 WT cells were treated either with NoA alone or co-treated with GCN2iB (10μM). Annexin V surface binding and Propidium Iodide uptake was measured after 48 and 72 h to assess viability of cells. Bar plots show mean ± SD of three biological replicates. Legend: AV−/PI− = live cells, AV+/PI− = early apoptosis, AV+/PI+ and AV−/PI+ = late apoptosis. Statistical significance was determined using a two-way ANOVA with Bonferroni correction for multiple comparisons, comparing percentages of live cells of corresponding concentrations between either WT and GCN2 KO cells or WT cells co-treated with GCN2iB at every time point. \**P* < 0.05, \*\**P* < 0.01, \*\*\**P* < 0.001, \*\*\*\**P* < 0.0001 f. HAP-1 WT and GCN1 KO clones were treated with 50 nM of NoA. Annexin V surface binding and Propidium Iodide uptake was measured after 48 and 72 h to assess cell viability. Bar plots show mean ± SD of three biological replicates. Legend: AV−/PI− = live cells, AV+/PI− = early apoptosis, AV+/PI+ and AV−/PI+ = late apoptosis. Statistical significance was determined using a two-way ANOVA with Bonferroni correction for multiple comparisons, comparing percentages of live cells of corresponding concentrations between WT and GCN1 KO clones. \*\**P* < 0.01, \*\*\**P* < 0.001 g. WB analysis comparing HAP-1 WT and ZAKα KO cells after treatment with 50 nM of NoA. Samples were collected at indicated timepoints and lysates were probed with the indicated antibodies. h. NB analysis of tRNA PRO charging status in HAP-1 cells. Cells were treated either with NoA or Halofuginone at indicated concentrations for 6 h.

To determine which ISR kinase mediates the response to NoA, we used HAP-1 KO cells lacking each of the four eIF2α kinases. Remarkably, only GCN2-deficient cells failed to induce ATF4 and its downstream effector CHOP following NoA treatment, indicating that GCN2 drives ISR activation under these conditions (Fig. 3b, supp. Fig. 3a). Notably, ATF4 and CHOP induction occurred independently of PERK, further supporting the notion that the ER is not the primary target of NoA.

To confirm that GCN2-dependent ISR activation is not restricted to HAP-1 cells, we examined eIF2α phosphorylation in additional cell lines using the GCN2 inhibitor GCN2iB. Co-treatment with GCN2iB abolished NoA-induced ISR signaling and downstream pro-apoptotic responses in HAP-1, Nalm-6, and HCT-116 cells, indicating conservation of this mechanism across different cell types (Fig. 3c).

We next performed time-resolved western blot analysis in HAP-1 WT and GCN2 KO cells to examine ISR activation kinetics (Fig. 3d). In WT cells, ATF4 was rapidly induced as early as 2 h after NoA treatment, whereas the pro-apoptotic effector CHOP was upregulated only after 12 h. In contrast, GCN2 KO cells failed to induce either ATF4 or CHOP. To determine whether GCN2-dependent ISR activation contributes functionally to mitochondrial apoptosis, we assessed cell viability following NoA treatment in the absence of GCN2, either genetically (GCN2 KO) or pharmacologically using the inhibitor GCN2iB. Annexin V/PI staining revealed that both GCN2-deficient cells and WT cells co-treated with GCN2iB showed significantly increased survival after NoA treatment (Fig. 3e, supp. Fig. 3b).

Recent studies have shown that GCN2 activation depends on recruitment of its binding partner GCN1 to collided ribosomes^37,38^. We therefore asked whether GCN1 is also required for ISR activation following NoA treatment. To test this, we compared cell survival of HAP-1 WT and GCN1 KO cells after NoA exposure and found that GCN1-deficient cells displayed a level of protection from cell death comparable to that observed in GCN2 KO cells, underscoring the importance of GCN1 for ISR activation (Fig. 3f). Consistently, western blot analysis confirmed that GCN1 is necessary for ATF4 upregulation in HAP-1 cells following NoA treatment (supp. Fig. 3c).

We next asked whether ISR induction coincides with activation of the ribotoxic stress response (RSR), which can also be triggered by ribosome collisions. The mitogen-activated protein kinase kinase kinase (MAPKKK) ZAKα is a key mediator of the RSR and can promote pro-apoptotic signaling through the JNK/JUN pathway during prolonged translational stress^38,39^. As TRRUST analysis revealed upregulation of c-JUN target genes in NoA-treated HAP-1 cells and NoA-induced apoptosis was partially dependent on the JNK target BIM (supp. Fig. 2g), we investigated whether ZAKα is activated upon NoA treatment. Time-resolved western blot analysis of NoA-treated HAP-1 WT cells showed rapid JNK phosphorylation as early as 1 h after treatment (Fig. 3g). This early phosphorylation was strictly ZAKα-dependent, as ZAKα KO cells displayed markedly reduced phospho-JNK at early time points, although the signal increased at later stages of treatment. To assess the functional contribution of ZAKα signaling to NoA-induced apoptosis, we compared survival of HAP-1 WT and ZAKα KO cells after treatment with graded concentrations of NoA. Annexin V/PI staining revealed increased survival of ZAKα KO cells after 72 h of treatment (Supplementary Fig. 3d).

Having established that NoA activates IRE1α and that ZAKα KO cells retain JNK phosphorylation at later time points, we investigated whether ZAKα and IRE1α both contribute to JNK activation. Time-resolved western blot analysis of NoA-treated HAP-1 WT and IRE1α KO cells revealed rapid JNK phosphorylation, whereas XBP1 splicing (a hallmark of IRE1α activation) occurred only at later time points (supp. Fig. 3e)^40^. In IRE1α KO cells, XBP1 splicing was completely abolished while early JNK phosphorylation remained intact. However, JNK phosphorylation failed to increase at later time points in the absence of IRE1α, suggesting that early JNK activation is primarily driven by ZAKα and subsequently reinforced by IRE1α signaling. Notably, among the UPR components examined, only loss of IRE1α conferred a measurable survival advantage following NoA treatment (supp. Fig. 3f). This protective effect was not observed in XBP1-deficient cells, indicating that it depends on the kinase module of IRE1α and its ability to promote JNK phosphorylation rather than on its splicing module (supp. Fig. 3g).

Finally, to assess whether NoA induced RSR signaling may be a consequence of cytosolic amino acid limitation, we analyzed tRNA charging by Northern blot analysis (NB). Uncharged tRNAs migrate faster than aminoacylated tRNAs, resulting in a detectable mobility shift when probed with radiolabeled probes specific for the corresponding tRNA species. Using a probe targeting tRNA Proline, we observed an increased fraction of uncharged tRNA Pro following NoA treatment (Fig. 3h). A similar shift was observed in cells treated with halofuginone, a prolyl-tRNA synthetase inhibitor used as a positive control. These results indicate that both NoA and halofuginone increase the fraction of uncharged tRNA Pro by a yet to be identified mechanism.

### Nostatin A binds and inhibits V-type H⁺-translocating ATPase

Next, we sought to identify the molecular target of NoA using a chemoproteomic competition assay^41^. NoA was immobilized on NHS-activated Sepharose beads via amidation of its N-terminal amino group, generating an affinity matrix (iNoA) to capture target proteins from cell lysates (supp. Fig. 4a). Pulldown competition experiments were performed using HAP-1 lysates in the presence of DMSO or increasing concentrations of soluble NoA. In this setup, free NoA competes with immobilized iNoA for binding to target proteins, resulting in dose-dependent depletion from the pulldown. Captured proteins were quantified by LC–MS/MS using a bottom-up proteomics workflow. Competition across nine concentrations enabled dose–response curve fitting and calculation of EC50 binding affinities for competed proteins^42^. Among the significantly and dose-dependently depleted proteins, several V-ATPase subunits and associated factors were identified (Fig. 4a). Visualization of protein–protein interactions using STRINGdb revealed a V-ATPase network, and GO analysis showed enrichment of terms related to lysosomal acidification (supp. Fig. 4b)^43^.

**Figure 4:**
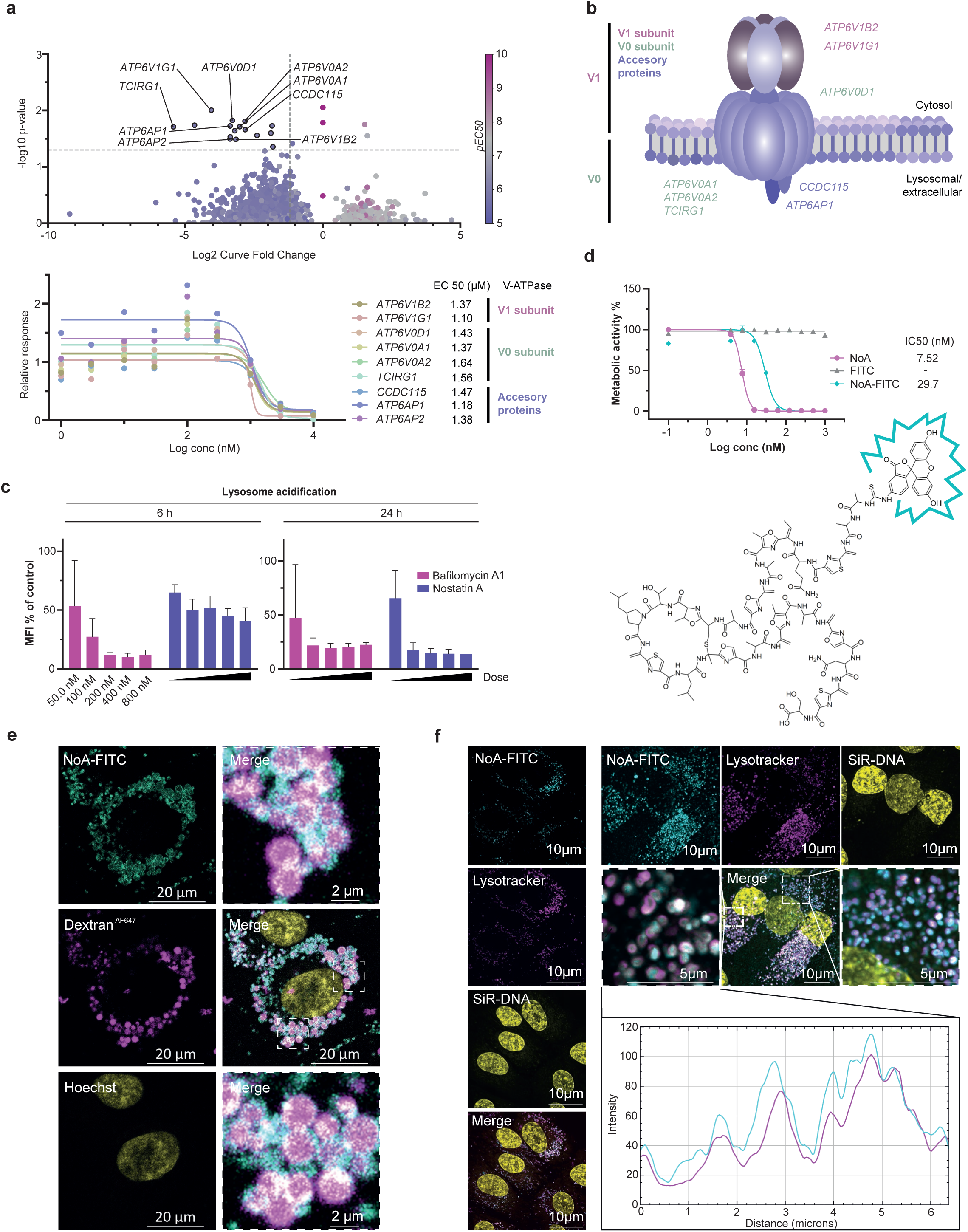
Nostatin A binds and inhibits V-type H⁺-translocating ATPase. a. Top: Overview of the chemoproteomic competition assay results. Volcano plot showing the significance of protein dose–response curves versus the log₂ fold change of the maximal effect size, calculated using the CurveCurator pipeline^42^. Proteins displaying significant dose–response behavior are highlighted. Bottom: Dose–response curves of the significant hits identified in the NoA chemoproteomic competition binding assay. Hits are annotated according to their corresponding V-ATPase subunit. b. Protein-protein interaction network of significantly depleted targets identified as NoA interactos. Individual proteins are represented as nodes, while predicted or established interactions between them are represented as edges. Network and GO term analysis were generated using the STRING database (version 12.0). c. HAP-1 WT cells were treated with NoA or BafA1 at 50, 100, 200, 400 and 800 nM for 6 or 24 h. Lysosomal acidification was assessed using the pH-sensitive dye LysoTracke. Fluorescence intensity was quantified by flow cytometry as mean fluorescence intensity (MFI) and normalized to the DMSO control d. Top: HCT-116 WT cells were treated with graded concentrations of NoA, NoA-FITC, or FITC for 72 h. Metabolic activity was assessed using the CellTiter-Glo® Luminescent Cell Viability Assay. IC₅₀ values are shown as mean ± SD of N = 2 biological replicates and were determined by linear regression. Bottom: Chemical structure of FITC-labeled NoA. e. RPE-1 cells were pre-treated with Alexa Fluor™ 647–labeled dextran to label endolysosomal compartments, followed by a wash step and treatment with 100 nM NoA-FITC. Images were acquired after 16 h of NoA-FITC treatment using confocal fluorescence microscopy to assess co-localization with dextran-labeled vesicles. NoA-FITC is shown in cyan, dextran in purple, and nuclei (Hoechst) in yellow. f. Super-resolution spinning disk confocal microscopy of RPE-1 cells treated with 100 nM NoA-FITC (cyan) for 6 h and co-stained with LysoTracke (purple) and the DNA dye SiR-DNA (yellow). Representative images for each channel are shown (left). Co-localization of NoA-FITC and LysoTracker was assessed by scanning representative frames and comparing fluorescence intensity profiles of the FITC and LysoTracker channels. Overlapping intensity peaks indicate spatial co-localization of NoA-FITC with lysosomes (right).

V-ATPases consist of two major components: the V1 and the V0 subcomplex (Fig. 4b)^44^. The cytosolic V_1_ subunit hydrolyzes ATP to subsequently enable H^+^ pumping through the transmembrane V_0_ domain. Indeed, we found multiple V_0_ and V_1_ subunits, such as ATP6V0A1, ATP6V0A2, ATP6V0D1, ATP6V1G1 and ATP6V1B2, all competed with overlapping dose response curves with EC50s in the range of 1-2 μM. We also found V-ATPase accessory proteins ATP6AP1 and ATP6AP2 to be outcompeted. These observations suggest that NoA binds to V-ATPase in HAP-1 cell lysate.

To functionally validate the chemoproteomics results, we assessed lysosomal acidification following NoA treatment and compared it with the bona fide V-ATPase inhibitor bafilomycin A1 (BafA1) using LysoTracker (Fig. 4c, supp. Fig. 4d)^45^. Both NoA and BafA1 caused a time-dependent decrease in LysoTracker signal, indicating increased lysosomal pH. Lysosomal pH was further examined using a pH-sensitive fluorescent dextran^AF647^ probe. In this assay, fluorescence is quenched in acidic lysosomes but increases upon V-ATPase inhibition. Treatment of MDA-MB231 breast cancer cells with the V-ATPase inhibitor concanamycin A (ConA) strongly increased fluorescence, validating the assay, and NoA produced a similar yet delayed response, consistent with lysosomal deacidification (supp. Fig. 4c). Together, these results support the notion that NoA inhibits V-ATPase function. The faster kinetics observed with BafA1 and ConA likely reflect their higher membrane permeability, whereas the size and polarity of NoA suggest limited diffusion across the plasma membrane, possibly requiring active uptake.

To determine whether NoA enters cells and localizes to V-ATPase–containing compartments, we generated a fluorescent probe by labeling NoA with fluorescein isothiocyanate (FITC) via direct conjugation, taking advantage of the free N-terminus being the only primary amine in the NoA molecule and resulting in FITC linked to NoA via thiourea bond (NoA-FITC) (Fig. 4d). We first confirmed that FITC-NoA retained biological activity, showing a modest reduction in killing potency compared with NoA in metabolic activity assays in HCT-116 WT cells. Using fluorescence microscopy, we then examined its cellular uptake and localization. FITC alone did not accumulate in intracellular compartments, whereas FITC-NoA formed distinct intracellular vesicular structures in RPE-1 cells, deemed more suitable for immunofluorescence (IF) microscopy analyses (supp. Fig. 4e). Confocal imaging of RPE1 cells revealed that NoA-FITC localized to the cytosol as discrete foci. Using SiR-DNA to mark the nucleus, we confirmed that NoA-FITC was entirely excluded from the nucleoplasm. (supp. Fig. 4f). Notably, extended incubation periods led to increased aggregation of these foci and the appearance of hollow, vesicle-like structures, indicating possible endosomal sequestration.

We then proceeded to perform an image-based time course analysis to determine the uptake kinetics of FITC-NoA (supp. Fig. 4g). NoA-FITC initially localized to the plasma membrane and steadily accumulated in cytosolic vesicles, with foci forming as early as 30 min after treatment. Considering the steady accumulation of NoA-FITC in these foci and the chemical properties of NoA (such as molecular weight and polarity), we reasoned that NoA/NoA-FITC is taken up by an active process such as endocytosis or micropinocytosis and steadily accumulating in lysosomes, where it inhibits V-ATPase function.

To validate this hypothesis, RPE-1 cells were pre-treated with fluorescently labeled dextran (Dextran^AF647^), followed by NoA-FITC treatment, and analyzed by live-cell fluorescence imaging to assess signal co-localization (Fig. 4e). Consistently, we observed co-localization of both signals, with dextran accumulating within the lumen of intracellular vesicles, while the NoA-FITC signal preferentially localized to the periphery of the same structures. This pattern is consistent with dextran uptake via fluid-phase endocytosis and membrane-associated localization of NoA-FITC.

To further confirm that these vesicles represent lysosomes and that NoA-FITC co-localizes with them, we performed live-cell fluorescence imaging of RPE-1 cells stained with LysoTracker following NoA-FITC treatment (Fig. 4f). High-resolution confocal microscopy equipped with a SoRa disk, revealed that LysoTracker signal intensity correlated with the NoA-FITC signal across multiple frames, indicating spatial co-localization. Notably, lysosomal acidification was not yet substantially impaired at the 6 h time point, as previously determined by flow cytometry, which likely explains the persistence of the LysoTracker signal under these conditions.

### V-ATPase inhibition activates the RSR/ISR and synergizes with BH3 mimetic treatment

Although several studies have reported cell death following V-ATPase inhibition, the underlying pro-apoptotic signaling pathways remain poorly characterized.^12^ We therefore asked whether the stress signaling cascade triggered by NoA could be generalized to other well-established V-ATPase inhibitors, as this may help to enable their potential therapeutic applications.

We first asked whether the established V-ATPase inhibitor BafA1 induces cell death with kinetics similar to NoA. Cell cycle profiling by flow cytometric DNA content analysis revealed that, like NoA, BafA1-treated HAP-1 WT cells initially underwent a G1 arrest before engaging apoptosis (Fig. 5a). Consistently, EdU incorporation assays showed a sharp decrease in S-phase activity following BafA1 treatment, further supporting the notion that V-ATPase inhibition induces an early G1 arrest (supp. Fig. 5a). BAX/BAK-deficient cells did not display an increase in the SubG1 population but also failed to re-enter the cell cycle after prolonged BafA1 exposure, mirroring our observations with NoA. We next examined whether BafA1 induces mitochondrial apoptosis. HAP-1 WT and BB dKO cells were treated with graded concentrations of BafA1, and viability was assessed by Annexin V/PI staining (Fig. 5b). Similar to NoA-treated cells, Annexin V binding was almost completely abolished in BB dKO cells, confirming that BafA1 induces BAX- and BAK-dependent apoptosis.

**Figure 5:**
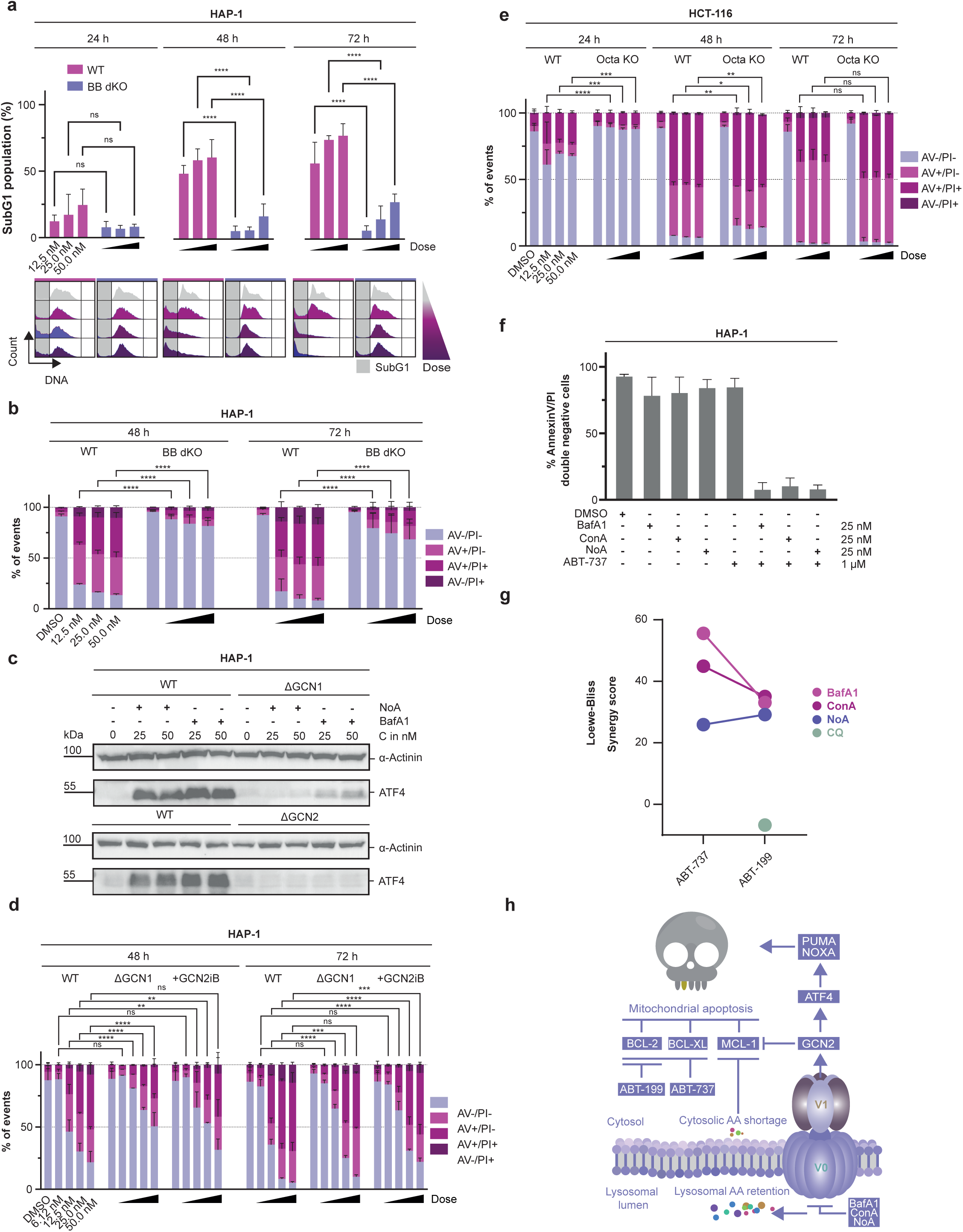
V-ATPase inhibition activates the RSR/ISR and synergizes with BH3 mimetic treatment. **a.** Bottom: Representative cell cycle profiles of HAP-1 WT and BB dKO cells after exposure to 12.5, 25 and 50 nM of NoA. Right: Quantification of the SubG1 population represented as bar graphs, showing means and SD of N=3 biological replicates. Statistical significance was determined using a two-way ANOVA with Bonferroni correction for multiple comparisons, comparing SubG1 percentages of corresponding concentrations of WT with BBdKO cells at every time point. \*\*\*\**P* < 0.0001 **b.** HAP-1 WT and HAP-1 BB dKO KO cells were treated with 12.5, 25 and 50 nM of BafA1. Annexin V surface binding and Propidium Iodide uptake was measured after 48 and 72 h to assess viability of the cells. Bar plots show mean ± SD of three biological replicates. Legend: AV−/PI− = live cells, AV+/PI− = early apoptosis, AV+/PI+ and AV−/PI+ = late apoptosis. Statistical significance was determined using a two-way ANOVA with Bonferroni correction for multiple comparisons, comparing percentages of live cells of corresponding concentrations between WT and BB dKO cells at every time point. \*\*\*\**P* < 0.0001. **c.** WB analysis of HAP-1 WT, GCN1 KO, and GCN2 KO cells treated with NoA or BafA1 (25 or 50 nM) for 24 h. Cell lysates were probed for the indicated proteins. **d.** HAP-1 WT and GCN1 KO cells were treated with BafA1 (6.25, 12.5, 25, or 50 nM). HAP-1 WT cells were treated either with BafA1 alone or in combination with the GCN2 inhibitor GCN2iB (10 μM). Annexin V surface binding and propidium iodide (PI) uptake were measured after 48 and 72 h to assess cell viability. Bar plots represent mean ± SD of three biological replicates. Legend: AV−/PI−, live cells; AV+/PI−, early apoptosis; AV+/PI+ and AV−/PI+, late apoptosis. Statistical significance was determined by two-way ANOVA with Bonferroni correction for multiple comparisons, comparing percentages of live cells at corresponding concentrations between WT and GCN1 KO cells or WT cells co-treated with GCN2iB at each time point. \*\**P* < 0.01, \*\*\**P* < 0.001, \*\**P* < 0.0001. **e.** HCT-116 WT and Octa KO clones were treated with 12.5, 25 and 5 0nM of BafA1. Annexin V surface binding and Propidium Iodide uptake was measured after 24, 48 and 72 h to assess cell viability. Bar plots show mean ± SD of three biological replicates. Legend: AV−/PI− = live cells, AV+/PI− = early apoptosis, AV+/PI+ and AV−/PI+ = late apoptosis. Statistical significance was determined using a two-way ANOVA with Bonferroni correction for multiple comparisons, comparing percentages of live cells of corresponding concentrations between WT and Octa KO clones. \**P* < 0.015, \*\**P* < 0.004, \*\*\**P* < 0.001, \*\*\*\**P* < 0.0001 **f.** HAP-1 WT cells were treated with indicated V-ATPase inhibitors (25 nM) or ABT-737 (1 μM) alone or in combination for 24 h. Viability was assessed by Annexin V surface binding and Propidium Iodide uptake. Viable fraction was defined as double negative cells and is depicted as mean ± SD of three biological replicates. **g.** Dot plot showing synergy scores for combinations of V-ATPase inhibitors with BH3 mimetics (ABT-737 and ABT-199). Synergy scores were calculated using the SynergyFinder web tool based on concentration matrices of two biological replicates. Cell viability was defined as the fraction of Annexin V−/PI− (double-negative) cells, representing live cells. **h.** Proposed mechanism for V-ATPase inhibition induced cell death

A key finding of our study is that V-ATPase inhibition by NoA activates the ISR through GCN1 and GCN2 and the RSR through ZAKα. As this link between V-ATPase inhibition and ISR activation has not been described previously, we asked whether the established V-ATPase inhibitor BafA1 would trigger a similar stress signaling profile. Hence, we first examined whether BafA1 activates the ISR via GCN1/2. HAP-1 WT, GCN1 KO, and GCN2 KO cells were treated with NoA or BafA1, and ATF4 induction was assessed by western blot (Fig. 5c). Consistent with our previous findings, ATF4 upregulation following V-ATPase inhibition strictly depended on the presence of GCN1 and GCN2. Moreover, GCN1-deficient cells, as well as WT cells co-treated with the GCN2 inhibitor GCN2iB, showed increased survival after BafA1 exposure, underscoring the contribution of chronic ISR activation to apoptosis following V-ATPase inhibition (Fig. 5d).

To determine whether ISR activation is a general consequence of lysosomal dysfunction or a specific feature of V-ATPase inhibition, we treated HAP-1 WT and GCN1 KO cells with three V-ATPase inhibitors (NoA, BafA1, and ConA) and two lysosomotropic agents (chloroquine (CQ) and monensin) (supp. Fig 5b). GCN1-dependent ATF4 induction was observed only with V-ATPase inhibitors. At the concentrations tested, chloroquine did not induce the ISR, while monensin triggered ATF4 upregulation independently of GCN1. These results highlight a shared stress signaling mechanism among V-ATPase inhibitors and distinguish their mode of action from that of lysosomotropic drugs. We next examined whether ATF4 induction following BafA1 treatment is conserved across different cell lines. Western blot analysis revealed robust ATF4 upregulation in HAP-1, Nalm-6, and HCT-116 cells upon BafA1 treatment (supp. Fig 5c).

Finally, we asked whether BafA1 also activates the RSR through ZAKα. HAP-1 WT and ZAKα KO cells were treated with BafA1 and JNK phosphorylation was assessed by western blot (supp. Fig. 5d). While WT cells showed a rapid increase in pJNK following BafA1 treatment, this signal was markedly reduced in ZAKα KO cells.

As we previously observed for NoA, V-ATPase inhibition–induced cell death was largely driven by depletion of pro-survival BCL-2 proteins rather than by upregulation of pro-apoptotic BH3-only proteins. To test whether this mechanism is conserved with BafA1, we used HCT-116 Octa KO cells and treated them with graded concentrations of BafA1 (Fig. 5e). Although cell death appeared mildly delayed in these cells compared with WT cells, no statistically significant difference in survival was observed at later time points. HCT-116 Octa KO cells also showed clear signs of caspase activation, as indicated by PARP cleavage, mirroring our observations following NoA treatment (Supplementary Fig. 5e). We next examined pro-survival BCL-2 family proteins in HAP-1, Nalm-6, and HCT-116 cells following BafA1 treatment to determine whether the effects observed with NoA were conserved. Consistent with our previous results, MCL-1 was the only pro-survival BCL-2 family member that was depleted, while BCL-2 and BCL-XL levels remained largely unchanged across cell lines (supp. Fig. 5f).

As V-ATPase inhibition did not induce rapid cell death but selectively reduced MCL-1 levels, we reasoned that treated cells may become increasingly dependent on the remaining pro-survival proteins BCL-2 and BCL-XL. We therefore hypothesized that simultaneous inhibition of these proteins using the BH3 mimetic ABT-737 would sensitize cells to V-ATPase inhibition^46^. To test this, HAP-1 WT cells were treated with the V-ATPase inhibitors NoA, BafA1, or ConA alone, or in combination with ABT-737, and viability was assessed by Annexin V/PI staining (Fig. 5f, supp. Fig. 5g). While single treatments caused little cell death after 24 h, the combination treatments markedly reduced cell viability. We next tested if the clinically approved BCL-2 inhibitor ABT-199 (venetoclax) would also produce a synergistic effect, as BCL-XL inhibition is linked to dose-limiting thrombocytopenia. Synergy scores calculated using the Loewe–Bliss model using SynergyFinder confirmed clear synergy for ABT-737 and ABT-199, consistent with increased BCL-2 dependency following V-ATPase inhibition (Fig. 5g)^47^. Notably, synergy was not observed when combining ABT-199 and the lysosomotropic agent chloroquine, confirming specificity of synergy for V-ATPase inhibitors.

## Discussion

In this study we aimed to identify the molecular target of the cyanobacterial natural product NoA. Through the initially target-agnostic characterization of NoA-induced cell death, we found that NoA leads to the chronic activation of the ISR via GCN2, followed by a cell cycle arrest and subsequent induction of mitochondrial apoptosis. NoA target identification through chemoproteomics showed, that this was a downstream consequence of V-ATPase inhibition. We established that GCN2-dependent ISR activation in response to V-ATPase inhibition is not a unique feature of NoA, but a conserved mechanism activated by different V-ATPase inhibitors. Through our cell death centric approach, we also discovered that V-ATPase inhibition leads to an increased dependency on the pro-survival proteins BCL-2 and BCL-XL, following MCL-1 depletion across multiple cancer cell lines. We propose that this dependency can be therapeutically leveraged through the combination of BH3 mimetics and V-ATPase inhibitors to enhance cancer cell killing, avoiding negative side effects linked to each class of compounds.

A major advance of this study is the identification of the lysosomal V-type H^+^-translocating ATPases as a candidate NoA interactor, suggesting that NoA may represent the first V-ATPase inhibitor described within the RiPP superfamily^8^. V-ATPases are evolutionarily conserved rotary proton pumps that are central to cellular bioenergetics and pH homeostasis^15^. In line with their fundamental role, they have repeatedly been identified as targets of natural products, including the plecomacrolides bafilomycin A1 and concanamycin A.

Using an unbiased chemoproteomic competition approach, we observed significant enrichment of multiple subunits and accessory proteins of the V-ATPase multi-protein complex. These findings strongly support an association between NoA and V-ATPases. However, while the chemoproteomic data are highly suggestive, they do not formally establish direct target engagement, and physical binding of NoA to individual subunits of the V-ATPase complex remains to be demonstrated. Importantly, however, our functional assays are fully consistent with vacuolar V-ATPase inhibition, as NoA-treated cells display hallmark consequences of impaired V-ATPase activity, including lysosomal dysfunction and loss of lysosomal acidification, mimicking the action of BafA or ConA^48,49^.

By fluorescently labeling NoA we were able to determine its subcellular location and spatiotemporal dynamics of uptake, establishing NoA’s MOA. Our data support a model in which the compound exerts its bioactivity through an endolysosomal trafficking–dependent mechanism. We show that the compound initially associates with the extracellular side of the plasma membrane, prior to internalization. Subsequent co-localization with fluorescently labeled dextran, a canonical marker of fluid-phase endocytosis and macropinocytosis, demonstrates that NoA is internalized through an endocytic or macropinocytotic route and accumulates over time in lysosomes^50^. We also demonstrate that NoA-FITC remains localized to the membrane throughout the time of uptake. This implies that the membrane integral V0 domain is likely the primary target of NoA. These observations support a model in which the peptide gains access to its target during its endocytic uptake and inhibits proton pumping into the endolysosomal lumen. This trafficking-dependent mode of action distinguishes the compound from small-molecule V-ATPase inhibitors such as BafA that freely diffuse across plasma membranes and suggests that intracellular routing is a critical determinant of its activity.

V-ATPases have emerged as attractive therapeutic targets in cancer, as many tumors rely on elevated autophagy and macropinocytosis to maintain amino acid supply and lysosomal function. In particular, K-Ras-driven tumors display higher rates of micropinocytosis that depend on V-ATPase recruitment to the plasma membrane^51,52^. However, clinical translation has been limited by toxicity and an incomplete understanding of how V-ATPase inhibition induces cell death, and which type of cell death modality is primarily engaged. As V-ATPase inhibition does not directly trigger lysosomal membrane permeabilization, leading to the release of cathepsins into the cytosol, or the activation of inflammasomes upon pathogen-induced lysosomal leakage^53^, our findings document for the first time how impaired lysosome function due to alkalinization drives the induction of apoptotic cell death (Fig. 5h).

Our data support a model where rapid activation of the ISR in a GCN1/GCN2–dependent manner, concomitant with ZAKα-dependent JNK phosphorylation and engagement of the RSR leads to *BAX/BAK* dependent mitochondrial cell death, a response likely enforced by the secondary activation of IRE1a, downstream of the ISR. Given that ribosome collisions activate GCN1/GCN2 and ZAKα, and that V-ATPase inhibition causes amino acid retention within lysosomes, these responses are consistent with cytosolic amino acid depletion, impaired tRNA charging and subsequent ribosomal collisions, eventually engaging the mitochondrial apoptosis pathway^20^. This observation is reminiscent of recent studies documenting that upon DNA damage, cleavage of a single tRNA by the transfer RNAase, SFLN11, suffices to trigger translational stalling, ribosomal collisions and BAX/BAK dependent apoptosis, independent of the tumor suppressor p53^54^.

Execution of mitochondrial apoptosis relies on BAX/BAK, usually activated downstream of BH3 only proteins that respond to different types of stressors, leading to inactivation of anti-apoptotic BCL-2 family members or direct activation of BAX/BAK^55^. Although ISR/RSR activation upon V-ATPase inhibition induces the expression of pro-apoptotic BCL-2 family members (PUMA, NOXA) via target gene transcription or JNK-mediated phosphorylation (BIM), genetic ablation of GCN1, GCN2, its chemical inhibition, or ZAKα deletion confer only limited protection from mitochondrial apoptosis. Similarly, IRE1a depletion or chemical inhibition also failed to provide substantial cell death protection, indicating a substantial degree of redundancy in apical cell death signaling^35^ and the need for depletion of anti-apoptotic proteins to engage apoptosis. Consistently, studying expression levels of pro-survival BCL-XL and MCL-1 and using cells lacking all pro-apoptotic BCL-2 proteins, we observed that cell death correlates with loss of pro-survival MCL-1. Notably, V-ATPase inhibition promotes NOXA-independent MCL-1 depletion, a finding of relevance, since MCL-1 is a known drug-resistance mechanism, in solid and hematological cancers^56,57^. Mechanistically, short-lived MCL-1 can decline via two distinct pathways after V-ATPase inhibition: pEIF2α dependent switch to cap-independent translation, and cytosolic amino acid depletion resulting from their lysosomal sequestration, reducing global translation efficiency^20,24^. The latter also counteracts the efficient transcriptional activation of BH3 only proteins. As such, MCL-1depletion appears critical to activate BAX/BAK without the need of efficient BH3 only protein induction. Consistently, studies from the Luo lab have shown that exogenous expression of BAX or BAK in cells lacking all BCL-2 family proteins suffices to induce mitochondrial apoptosis and caspase activation^35^. Consistently, we observed near normal caspase activation levels upon V-ATPase inhibition in HCT-116 cells devoid of all apoptosis-inducing BH3-only proteins (Octa KO), but not in cells lacking BAX/BAK, or those devoid of all major BCL-2 family members (BCL-2 all KO).

Notably, the transcriptional activation of BH3 only proteins PUMA and NOXA in response to V-ATPase inhibition appears to be a rather late event, driven in response to IRE1a activation in a secondary UPR-like response, mounted in response to a late-stage accumulation of unfolded proteins, indicated by TPE-MI activity 24h after treatment. This second signaling wave galvanizing a continued JNK and ATF4 response likely enforces the apoptotic response by transcriptional activation and/or posttranslational activation of BH3 only proteins, known to execute apoptosis in response to prolonged activation of the UPR^58,59^. However, the observation that none of these proteins appear to be rate-limiting for cell death yet cell death still proceeds in a BAX/BAK dependent manner opens up an exciting therapeutic opportunity. Most if not all currently used anti-cancer therapies rely on the activation of BH3 only proteins, which is frequently impaired by loss of its transcriptional regulators (e.g., p53), gene loss, or epigenetic silencing^55,60^. V-ATPase inhibition appears to be able to short-cut the need for efficient BH3-only protein engagement and the notion that high levels of synergy can be obtained by combination with BH3-mimetics cannot be emphasized enough. Our observation that selective inhibition of BCL-2 using venetoclax, the only clinically approved BH3 mimetic, synergizes with V-ATPase inhibition opens the door to apply tolerable doses of V-ATPase inhibitors to patients, but, maybe even more importantly, the noted depletion of MCL-1, may pave a way for rendering solid cancers BCL-2 dependent^61^. This strategy is highly attractive, as direct inhibition of MCL-1 or BCL-XL in cancer patients suffered repeat set-backs due to cardiac toxicity or thrombocytopenia, respectively^62–65^.

Together, our data establish MCL-1 depletion as a critical downstream event of V-ATPase inhibition and provide a mechanistic framework linking V-ATPase inhibition to mitochondrial apoptosis. These findings help explain the long-recognized cytotoxicity associated with V-ATPase inhibitors and warrant caution in the use of agents such as bafilomycin A1 in studies of autophagy. At the same time, the selective loss of MCL-1 creates a therapeutically exploitable vulnerability by rendering cancer cells increasingly dependent on the remaining pro-survival BCL-2 family members. In this context, combining V-ATPase inhibitors with BH3 mimetics targeting BCL-2 or BCL-XL may represent a strategy to enhance cancer cell killing while potentially allowing the use of lower, more tolerable doses of V-ATPase inhibitors.

## Materials and Methods

### Cell culture and drug treatment

Human HAP-1 (WT and indicated knock-out clones) cells were obtained from Horizon Genomics and cultured in Iscove’s Modified Dulbecco’s Medium (IMDM) (I6529, Sigma-Aldrich). Nalm-6 cells were grown in RPMI-1640 (R8758, Sigma-Aldrich) and BAX/BAK or caspase-deficient mutants used here were characterized before^27,35^. HCT-116 cells were maintained in Dulbecco’s modified Eagle’s medium (DMEM) (D5796, Sigma-Aldrich). Human retinal pigment epithelial cells (RPE-1) were maintained in DMEM. Growth media were all supplemented with supplemented with 10% fetal bovine serum (FBS) (F7524, Sigma-Aldrich), penicillin (100 U), and streptomycin (0.1 mg/ml) (P4333, Sigma-Aldrich). Trypsin-EDTA solution (T4049, Sigma-Aldrich) was used to detach adherent cells from dishes. All cells were grown at 5% CO_2_ and 37°C. Cells used for this study were regularly checked for mycoplasma contamination by PCR.

Compounds use in this study include: Q-VD-OPH (CAY15260-1, Cayman Chemicals), used at a concentration of 10 μM; GCN2iB (HY-112654, MedChemExpress) used at a concentration of 10 μM; ABT-737 (HY-50907, MedChemExpress) and ABT-199 (HY-15531, MedChemExpress) used at indicated concentrations

### Generation of isogenic knock out cell lines using CRISPR-CasG

Isogenic KO cell lines in HAP-1, Nalm-6 and HCT-116 (if not published previously) were generated using lentiviral delivery of Cas9 and the sgRNA using the lentiCRISPR V2 backbone (#52961, Addgene)^66^.sgRNA sequences were designed with VBCscore and ordered at Microsynth Austria. They were subsequently cloned into the lentiCRISPR V2 backbone, following the protocol provided by Addgene. Whether the sgRNA was cloned correctly into the vector backbone was confirmed by Sanger sequencing (Microsynth Austria).

HEK293T cells were used for virus production. Cells were cotransfected with sgRNA containing lentiCRISPR V2 and viral packaging genes containing vectors pSPAX2 and VSVg, using polyethylenimine (PEI; 23966-100, Polysciences Europe GmbH) in a 1 μg:2 μl DNA:PEI ratio. HEK293T cells were then left for 24h before medium was changed to fresh DMEM complete. Medium was harvested after 72h and filtered using 0.45-μm polyethersulfone syringe filters to remove cellular debris. Virus was either used directly for transduction or stored at -80°C.

Cells of interest were seeded 24h before transduction. Transduction was performed by adding viral supernatant at a volume of either 50 or 25% of the total volume that cells were seeded in. Cells were incubated with the viral supernatant for 48h before transduced cells were selected using puromycin (13884, Cayman Chemical) at a concentration of 1 μg/ml. Double knock-out cell lines were generated through cotransduction of with two different viral supernatants at a 1:1 ratio. Selection conditions were maintained, till cells in the untransduced control were no longer viable. Polyclonal pools that survived selection were then expanded and knock-out efficiency was determined by western blot analysis.

Polyclonal pools were expanded, and the gene KO efficiency was tested by Western blotting. Polyclonal pools showing the highest reduction of signal on the protein level were then expanded and single cell clones were obtained through limiting dilution steps in a 96-well plate format. Wells containing only one colony were expanded and knock down efficiency was determined using western blot.

### Cell viability and metabolic activity assays

Cells were seeded in 96-well plates at a density of 5000 cells per well, incubated for 24 hours, and subsequently treated with indicated compounds or DMSO as vehicle control for 72h. Plates were then measured on the Plate Reader Platform Victor X3 model 2030 (PerkinElmer). Adenosine triphosphate (ATP) levels were measured as a proxy for viability using the luminescence based CellTiter-Glo assay (Promega G7573), following the manufacturer’s protocol. The experiments were performed in biological triplicates with three technical triplicates per condition. Where indicated, cells were co-treated with indicated compounds

The assays were analyzed using Prism version 10.0.3 (GraphPad). After background (medium only) subtraction, treatment conditions were normalized to the vehicle controls. Dose-response curves were calculated using GraphPad’s nonlinear regression curve fitting function, which then calculated the median inhibitory concentration (IC50).

### Western Blot

Cells seeded for western blot analysis were incubated for 24h before being treated with any compound. Cells were then harvested at relevant timepoints and washed twice with PBS before cell lysis. Pellets were resuspended in RIPA buffer [50 mM tris-HCl (pH 8), 150 mM NaCl, 1% NP-40, 0.5% sodium deoxycholate, 0.1% SDS, supplemented with 1 tablet/10 ml of cOmplete protease inhibitor, EDTA-free (4693132001, Roche), 1 tablet/10 ml of PhosSTOP™, phosphatase inhibitor (4906845001 Roche)] and lysed on a spinning wheel at 4°C for 60 min. Lysates were then spun down (10 min, 4°C at 12.000g) and protein concentration was assessed using a bicinchoninic acid assay (BCA) kit (Pierce™ BCA Protein Assay Kits, 23225, Thermo Scientific™). SDS-polyacrylamide gel electrophoresis was performed using 30 μg of sample in tris/glycine/SDS buffer. Proteins were then transferred on a nitro cellulose membrane (Amersham™ Protran® Western blotting membranes, nitrocellulose, 10600002, Cytiva™) in wet conditions tris/glycine buffer added with 20% ethanol. Membranes were blocked with 5% milk (Powdered milk, T145.3, Carl Roth) in Tris-buffered saline with 0.1% Tween® 20 detergent (TBS-T) for 1h. Primary antibodies were diluted in 1% milk with TBS-T and incubated with the primary antibody overnight at 4°C. Membranes were washed three times in TBS-T for 10 min and subsequently incubated with the corresponding secondary antibody for 1h at room temperature. Immunoreactive bands were detected by incubating membranes with ECL™ Select Western Blotting Detection Reagent (RPN2235, Cytiva™) for 3 min before fluorescent signals were detected on a ChemidocMP (Bio-Rad).

Antibodies were commercially obtained: PARP (#9542, Cell Signaling Technology), GAPDH (#2118, Cell Signaling Technology), β-Tubulin (sc-53140, Santa Cruz Biotechnology), phospho-H2A.X (07-164, Merck), anti-mouse (#7076, Cell Signaling Technology), anti-rat (#7077, Cell Signaling Technology), Caspase-3 (#9662,Cell Signaling Technology), XBP-1s (#12782, Cell Signaling Technology), ATF-6 (#65880, Cell Signaling Technology), α-Actinin (sc-17829, Santa Cruz), ATF-4 (#11815, Cell Signaling Technology), PERK (#3192, Cell Signaling Technology), eIF2α (#9722, Cell Signaling Technology), phospho-eIF2α (Ser51) (#9721, Cell Signaling Technology), GCN2 (#3302, Cell Signaling Technology), IRE1α (#3294,Cell Signaling Technology), Phospho-SAPK/JNK (Thr183/Tyr185)( #4668, Cell Signaling Technology), SAPK/JNK (#9252, Cell Signaling Technology), GAPDH (#2118, Cell Signaling Technology), CHOP (#2895, Cell Signaling Technology), BIM (#2933, Cell Signaling Technology), ZAKα (#BEYA301-993A-T, Szabo Scandic), PUMA (#4976 Cell Signaling Technology), NOXA (#14766, Cell Signaling Technology), BAX (#2772, Cell Signaling Technology), BAK(#3814, Cell Signaling Technology), MCL-1(#5452, Cell Signaling Technology); BCL-XL (#2762, Cell Signaling Technology); Protein markers used were either Broad Range Prestained Protein Marker (PL00002, Proteintech) or PageRuler^TM^ Plus (26619, Thermo Scientific).

### Northern Blot

Total RNA was extracted under acidic conditions using TRIzol. For electrophoresis, 6 μg of RNA were resolved on 8% polyacrylamide gels containing 7 M urea and supplemented with 20 μg/mL APM ([(N-acryloylamino)phenyl]mercuric chloride LGC Standards). RNA was subsequently transferred onto positively charged nylon membranes (Hybond-N, Amersham) using a semi-dry transfer system and cross-linked by UV irradiation. Membranes were pre-hybridized for 1 h at 40°C with rotation in 6× SSC buffer containing 10× Denhardt’s solution, 0.5% DMSO, and 100 μg/mL fish sperm DNA (Merck). Hybridization was performed overnight using radioactively labeled oligonucleotides ([γ-³²P] ATP). Following hybridization, membranes were washed with SSC buffers of decreasing concentration, wrapped in plastic, and exposed to a storage phosphor screen overnight. Signal detection was carried out using an Amersham Typhoon laser scanner (Cytiva). Oligonucleotide sequence for human tRNA-PRO^AGG^: GGGCTCGTCCGGGAT

### Cell cycle profiling and SubG1 assay

Cells were harvested at relevant timepoints and washed twice with Dulbecco Phosphate-Buffered Saline (PBS) (D8537, Sigma-Aldrich) before they were fixed in 70% cold EtOH and stored at -20°C for at least 6 hours. Fixed cells were then spun down at 900g for 4 min at room temperature and washed once with PBS before stained with a DNA staining solution [(propidium iodide (PI) (10 μg/ml) (14289, Cayman Chemical) and ribonuclease (RNase) A (100 μg/ml) (10109169001, Roche) in PBS]. The volume of the DNA staining solution was adjusted to not exceed the saturation point of 1 × 10^6^ cells/ml. Cells were stained for 30 min in the dark before being acquired on LSR Fortessa Cell Analyzer (BD Biosciences). FCS files exported from the LSR Fortessa Cell Analyzer were then analyzed using FlowJo v10.10 (FlowJo LLC). Gates were set to exclude doublets and the cell cycle profile was then displayed as a histogram in the PE channel. DMSO treated samples were used to determine the SubG1 threshold. Individual plots were exported as PNG files and assembled into summary plots in Adobe Illustrator version 29.1 (Adobe).

### Annexin V-PI staining

Cells were harvested at relevant timepoints and washed twice with PBS. The supernatant was aspirated and the cell pellet was resuspended in 100μl Annexin V Binding Buffer (422201, BioLegend). Samples were then stained with 5 μl of fluorescein isothiocyanate (FITC) Annexin V antibody (640906, BioLegend) and 10 μl of PI (0.5 mg/ml). Stained samples were mixed by pipetting up and down before they were incubated in the dark for 15 min at room temperature. After incubation, 400μl of Annexin V Binding Buffer was added to each sample. Samples were then acquired on an LSR Fortessa Cell Analyzer (BD Biosciences). FCS files were analyzed using FlowJo v10.10 (FlowJo LLC). Summary plots and statistical analysis of the data was performed in Prism version 10.0.3 (GraphPad).

### FACS based lysotracker readout

Cells were harvested and washed once with PBS before incubation with LysoTracker™ in PBS at a final concentration of 100 nM for 30 min at 37 °C. Following staining, cells were washed twice with PBS and immediately analyzed by flow cytometry using the LSR Fortessa Cell Analyzer (BD Biosciences). Mean fluorescence intensity (MFI) was used as a measure of lysosomal acidification. FCS files exported from the LSR Fortessa Cell Analyzer were then analyzed using FlowJo v10.10 (FlowJo LLC). DMSO-treated cells served as the control condition and were normalized to 100%, and relative MFI values were compared between treatment groups.

### Dextran-FITC lysosomal pH assay

Cells were seeded in 6-well plates one day prior to the experiment to reach ∼70% confluency. The following day, cells were incubated with FITC–dextran (MW 70,000; 100 µg/mL) for 24 h to allow uptake into endolysosomal compartments, as previously described^67,68^. After loading, cells were treated with Nostatin A (NoA, 100 nM), the lysosomotropic V-ATPase inhibitor concanamycin A (50 nM), or vehicle control. At the indicated time points, cells were washed with PBS, harvested by trypsinization, and analyzed by flow cytometry (BD Influx).

### EdU incorporation assay

To determine Edu incorporation we used the commercially available Click-iT^TM^ Plus EdU Alexa Fluor^TM^ 647 Flow Cytometry Assay Kit (C10634, Thermo Fisher Scientific). Experiments were performed according to the manufacturer’s instructions.

### TPE-MI staining

Cells were seeded in 12-well plates one day prior to the experiment to reach ∼70% confluency. After treatment cells were harvested and washed twice with PBS and spun down. Cell pellets were resuspended in 50 µL PBS containing 50 µM TPE-MI (a kind gift from Y. Hong) and incubated for 30 min at 37 °C^36^. After incubation, cells were washed with PBS and analyzed by flow cytometry using an LSR Fortessa Cell Analyzer (BD Biosciences). Flow cytometry data (FCS files) were analyzed were then analyzed using FlowJo v10.10 (FlowJo LLC), and statistical analyses were performed using Prism version 10.0.3 (GraphPad).

### Drug synergy analysis

Drug synergy was assessed using concentration matrix experiments in which cells were treated with increasing concentrations of the indicated V-ATPase inhibitor in combination with either ABT-737 or ABT-199 at different doses. After 24 h treatment, cell viability was determined by Annexin V/PI staining, and the fraction of Annexin V/PI double negative cells was defined as the viable population. Synergy scores were calculated from the resulting dose–response matrices using the SynergyFinder web application (Version 3.0) based on the Loewe–Bliss model^47^.

### Transcriptomics and TRRUST analysis

Poly-A mRNA was isolated with the NEBNext® Poly(A) mRNA Magnetic Isolation Module (E7490)from 500ng of total RNA. Enriched poly-A mRNA sequencing libraries were prepared using theNEBNext® Ultra™ II Directional RNA Library Prep Kit for Illumina (NEB). The fragment size ofthe libraries was assessed using the NGS HS analysis kit and a fragment analyzer system(Agilent). The library concentrations were quantified with a KAPA Kit (Roche). Libraries were sequenced as SR100 on a NovaSeq 6000 (SP lane) (Illumina, San Diego, California, USA). Basecalling was performed using Real-Time Analysis (RTA) BCL files were converted todemultiplexed fastq files with bcl2fastq v2.20.0.422. The quality of fastq files was checked with fastqc 0.11.9.. TRRUST transcription factor enrichment analysis was implemented by using Metascape (metascape.org). the top 200 upregulated genes were used for the TRRUST analysis.

### Isolation of NostatinA for biological assays

For each batch isolation approx. 10 g of cyanobacterium *Nostoc* sp. CCALA 1144 lyophilized biomass was ground in a mortar and extracted with 700 mL of 60% acetonitrile (ACN). The extraction was assisted by ultrasound treatment (10 minutes every 30 minutes) for a total duration of 3 hours. The resulting extract was centrifuged, and the supernatant was collected and evaporated to a small volume (approximately 1–5 mL). The remaining pellet was re-extracted with 500 mL of 100% methanol (MeOH) following the same procedure. Both extracts were analysed for the presence of the target compound. The combined concentrated supernatants were then diluted with 50% methanol to a final volume of 20–30 mL, ensuring that no precipitates were formed. The first purification was performed using a preparative HPLC system (Agilent 1200) equipped with a semipreparative YMC-Triart C18 column (250 × 10.0 mm, S-5 μm, 12 nm). The mobile phases were water with 0.1% acetic acid (solvent A) and acetonitrile (solvent B). The following gradient was applied: 0 min – 40% B; 5 min – 47% B; 35 min – 50% B; 37 min – 100% B; 45 min – 100% B; 46 min – 40% B, with a flow rate of 4 mL/min. The retention time was largely affected by concentration of co-eluting chlorophyll and its degradation products. Due to this the gradient between 47% and 50% B was slightly adjusted in each extraction batch to achieve optimal UV absorption peak separation. During the isolation process, this gradient range was modified between 47–50% up to 53–56% B. Since Nostatin A tends to decompose into a hydrated form in acidified solvents, concentration was performed using a C18 solid-phase extraction (SPE) column. The HPLC fraction was diluted with water to reduce the organic solvent concentration below 10% and then loaded onto the SPE column. After loading, the column was thoroughly washed with 100 mL of water, and Nostatin A was eluted with 100% methanol, yielding a non-acidified fraction that was evaporated to dryness using a rotary evaporator. Under optimal conditions, the first purification step yielded 2–5 mg of >90% pure Nostatin A, visible strong green/yellow pigment contamination remained. A second purification step was conducted on a semipreparative μBondapak Phenyl column (Prep 10 μm, 125 Å, 7.8 × 300 mm) using methanol (solvent B) and water (solvent A). The applied gradient was as follows: 0 min – 30% B; 9 min – 51% B; 58 min – 57% B; 58.1 min – 100% B; 65 min – 100% B; 65.1 min – 30% B; 76 min – 30% B, at a flow rate of 4 mL/min. As in the first purification, the main gradient was optimized for each batch, ranging between 40–50% and 54–58% B. The collected fractions were directly evaporated using a rotary evaporator. Under optimized conditions, the second purification yielded 1–2 mg of a white, highly purified Nostatin A (>98%).

### Preparation of the Nostatin A-Fluorescein isothiocyanate (FITC) conjugate

FITC was dissolved in 100 mM sodium carbonate buffer (pH 9.0) to prepare a stock solution at a concentration of 2 mg/mL. An aliquot of 100 µL of this FITC solution was added to a vial containing 1.25 mg of Nostatin A (dissolved in 0.5 mL of MeOH). The reaction mixture was incubated in the dark for 2 hours with gentle mixing every 30 minutes. The reaction was quenched by the addition of 100 µL of an ammonium chloride solution (1 mg/mL). The solvent was subsequently removed under reduced pressure using a rotary evaporator until dryness. The resulting residue was reconstituted in 50% methanol and subjected to microfractionation using a Thermo Dionex Ultimate 3000+ system equipped with an Arion C18 BIO300 Å column (3 µm, 100 mm × 2.1 mm). Chromatographic separation was performed using acetonitrile (solvent A) and water (solvent B), both containing 0.1% formic acid. The following gradient program was applied: 0–1 min, 15% A; 1–20 min, linear increase to 100% A; 20–25 min, 100% A; 25–30 min, decrease to 15% A; and 30–33 min, 15% A. The flow rate was maintained at 0.6 mL/min. Fractions were collected manually every 15 seconds and subsequently analysed for the presence of the desired labelled compound.

### NoA-FITC high-resolution imaging

RPE1 cells were seeded in Ibidi μ-Slide 8 Well glass-bottom dishes (Cat. no. 80827) at a density of 100,000 cells per well. The following day, cells were stained with 100 nM LysoTracker™ Red DND-99 (Invitrogen, Ref. 7528, Lot 1971192), 100 nM SiR-DNA (Spirochrome, 001_20.01, Lot 2401), or treated with 100 nM NoA-FITC.

Images were acquired using an Olympus IXplore SpinSR microscope equipped with a 60x silicone oil objective (NA 1.3) and a SoRa Super-Resolution Spinning Disk system, providing a resolution of ∼150 nm. Image acquisition was performed using *cellSens Dimensions* software, and images were processed via constrained iterative deconvolution for restoration.

A Z-stack was acquired with a step size of 0.3 μm. Excitation/emission settings were as follows: LysoTracker Red: excitation 561 nm, FITC-NoA compound: excitation 488 nm SiR-DNA: excitation 641 nm.

### NoA-FITC Opera High-Content Imaging

RPE1 cells were seeded in PhenoPlate™ 96-well microplates (Revvity, Cat. no. 6055302) at a density of 10,000 cells per well. The following day, cells were treated with 100 nM of NoA-FITC alone or in combination with SiR-DNA (100nM). Images were acquired at 0, 2, 4, and 6 h post-treatment. For long-term live-cell experiments, cells were maintained in a humidified Opera Phenix™ High-Content Screening System equipped with temperature and CO₂ control, and images were acquired at the indicated timepoints. Imaging was performed with a 40x water-immersion objective. Excitation/emission settings were: 488 nm excitation, emission 500-530nm, 561 nm excitation, emission 570-630 nm The acquired images were analyzed using *ImageJ* software, including background subtraction, fluorescence intensity measurements, and quantitative signal analysis.

To analyze the spatial correlation between the two fluorescence channels, intensity line profiles were generated from composite images using the RGB Profiler plugin in ImageJ/Fiji (National Institutes of Health, USA).

### Fluorescent dextran and NoA-FITC co-localization by confocal microscopy

RPE-1 cells were seeded at 5,000 cells per well in 200 μL DMEM in 96-well glass-bottom imaging plates (black walls, 0.2 mm glass bottom, half-area) and allowed to adhere for 24 h at 37 °C and 5% CO₂. For lysosomal labeling, cells were incubated with Alexa Fluor 647–conjugated dextran (D22914, Invitrogen, 10 kDa) at a final concentration of 0.05 mg mL⁻¹ for 1 h, followed by two washes with PBS. Cells were then treated with NoA–FITC (800 nM). Prior to imaging, cells were washed twice with PBS and stained with Hoechst 33342 (Invitrogen) for 5 min, followed by a PBS wash and resuspension in culturing medium. Imaging was performed 3, 8, 16, and 20 h after NoA–FITC treatment. To avoid cellular stress associated with repeated imaging, separate wells were treated and analyzed for each time point.

Fluorescence imaging was performed using a laser scanning confocal microscope (LSM 880, Zeiss) equipped with a temperature-controlled chamber maintained at 37 °C and a 63×/1.4 NA oil-immersion objective. After image acquisition, 96-well glass-bottom imaging plates were returned to the incubator under identical culture conditions until the next time point.

Fluorophores were excited sequentially with 405 nm (Hoechst), 488 nm (NoA–FITC) and 633 nm (Alexa Fluor 647) laser lines. Emission was detected at 427–481 nm (Hoechst), 508–561 nm (FITC) and 642–695 nm (Alexa Fluor 647). Fluorescence signals were detected using GaAsP photomultiplier detectors. Images were acquired in frame mode using ZEN Blue software (version 3.3). Visualization of acquired pictures was processed in the GIMP software (Version 3.0.8. Community, Free Software).

### Preparation of the iNoA pulldown affinity matrix

To immobilize Nostatin A to a solid phase matrix via its N-terminal amine, Nostatin A (1 µM) was incubated with DMSO-washed NHS-activated Sepharose beads (1 ml; ∼20 µM NHS groups per ml beads) and triethylamine (20 µl) in DMSO (2 ml) on an end-over-end shaker overnight at room temperature in the dark. Aminoethanol (50 µl) was then added to quench remaining NHS-activated carboxylic acid groups. After 16 h, the beads were washed with 10 ml DMSO followed by 30 ml ethanol to yield the iNoA affinity matrix for pulldown experiments.

### Preparation of HAP-1 lysates for chemoproteomic competition assays

HAP-1 cells were grown on 15 cm cell culture dishes in IMDM (see section ‘Cell culture and drug treatment’). Cells were washed twice with 10 mL of PBS and lysed via scraping in buffer containing 0.8% Igepal, 50 mM Tris-HCl pH 7.5, 5% glycerol, 1.5 mM MgCl₂, 150 mM NaCl, 1 mM Na₃VO₄, 25 mM NaF and 1 mM dithiothreitol (DTT), supplemented with protease inhibitors. After one freeze-thaw cycle lysates were cleared by centrifugation at 21,000g for 30 min at 4 °C. Protein concentration was determined by BCA assay and adjusted to 5 mg ml⁻¹ and 0.4% Igepal by dilution with detergent-free lysis buffer and buffer containing 0.4% Igepal, respectively.

### Chemoproteomic competition pulldown assay

Aliquots of lysate (0.5 ml) were pre-incubated with nine concentrations of free Nostatin A (1, 3, 10, 30, 100, 300, 1,000, 3,000, and 10,000 nM) or DMSO vehicle control for 1 h at 4 °C with gentle shaking. Samples were then transferred to 17 µl of affinity matrix equilibrated in lysis buffer in a filter plate (Porvair Sciences, #240002) and incubated for 30 min at 4 °C with gentle shaking. Beads were washed once with 1 ml lysis buffer containing 0.4% Igepal and twice with 2 ml lysis buffer containing 0.2% Igepal. Filter plates were centrifuged at 300 g for 2 min to remove residual detergent-containing buffer and washed three additional times with detergent-free lysis buffer. Pulled down proteins were denatured on-bead in 40 µl 8 M urea in Tris-HCl containing 10 mM dithiothreitol (DTT) for 30 min at 37 °C with shaking at 700 rpm. Iodoacetamide (4 µl of 550 mM) was added for alkylation at 37 °C with shaking at 700 rpm. Samples were diluted to 1 M urea with 250 µl 40 mM Tris-HCl and digested overnight at 37 °C with trypsin (final concentration 1 ng ml⁻¹). Peptides were recovered, acidified with 10 µl 10% formic acid, desalted using C18 StageTips, vacuum-dried and stored at −20 °C until LC–MS/MS analysis.

### LC-MS/MS data acquisition of competition pulldown assay

For proteomic data acquisition, a nanoflow LC−ESI-MS/MS setup comprised of a Dionex Ultimate 3000 RSLCnano system coupled to a Fusion Lumos mass spectrometer (both ThermoFisher Scientific Inc.) was used in positive ionization mode. MS data acquisition was performed in data-dependent acquisition (DDA) mode. For proteome analyses, half of the competition pulldown peptides were delivered to a trap column (Acclaim™ PepMap™ 100 C18, 3 μm, 5 × 0.3 mm, Thermo Fisher Scientific) at a flowrate of 5 μL/min in HPLC grade water with 0.1% (v/v) TFA. After 10 min of loading, peptides were transferred to an analytical column (ReproSil Pur C18-AQ, 3 μm, Dr. Maisch, 500 mm × 75 μm, self-packed) and separated using a stepped gradient from minute 11 at 4% solvent B (0.4% (v/v) FA in 90% ACN) to minute 61 at 24% solvent B and minute 81 at 36% solvent B at 300 nL/min flow rate. The nano-LC solvent A was 0.4% (v/v) FA HPLC-grade water. MS1 spectra were recorded at a resolution of 60,000 using an automatic gain control target value of 4 × 105 and a maximum injection time of 50 ms. The cycle time was set to 2 seconds. Only precursors with charge state 2 to 6 which fall in a mass range between 360 to 1300 Da were selected and dynamic exclusion of 30 s was enabled. Peptide fragmentation was performed using higher energy collision dissociation (HCD) and a normalized collision energy of 30%. The precursor isolation window width was set to 1.3 m/z. MS2 spectra were acquired at a resolution of 30,000 with an automatic gain control target value of 5 × 104 and a maximum injection time of 54 ms.

### Chemoproteomic protein identification and quantification

Protein identification and quantification was performed using MaxQuant (v2.4.9.0) by searching the LC–MS/MS data against all canonical protein sequences as annotated in the Uniprot reference database (uniprotkb_reviewed_true_canon_human, 20,434 entries, downloaded 29 April 2024) using the embedded search engine Andromeda^69^. Carbamidomethylated cysteine was set as fixed modification and oxidation of methionine and amino-terminal protein acetylation as variable modifications. Trypsin/P was specified as the proteolytic enzyme, and up to two missed cleavage sites were allowed. The minimum length of amino acids was set to seven, and all data were adjusted to 1% peptide spectrum matches and 1% protein false discovery rate. Label-free quantification and match between runs was enabled^69^.

### Competition pulldown assay data analysis

We employed the CurveCurator pipeline to calculate the dose-dependent residual binding of each protein to the affinity matrix relative to the DMSO control and to estimate effect potency (EC50), effect size, and the statistical significance of the observed dose response^42^. MaxQuant Protein intensities were used as input for relative protein abundance.

## Data availability

The LC-MS/MS chemoproteomics data, including the used protein reference database has been deposited in the MassIVE proteomics database with the dataset identifier **MSV000100G35**.

## Acknowledgements

We thank the Proteomics team of the Molecular Discovery Platform at CeMM for mass-spectrometric data acquisition and the Vienna Biocenter NGS Core Facility or expert assistance. We thank the Kubicek lab for providing Horizon Discovery HAP-1 KO cell lines. AV and PH acknowledge support by the Czech Science Foundation and Austrian Science Fund (Projects # 21-05649K and 10.55776/I5133). CeMM is supported by the Austrian Academy of Sciences (OeAW); AITHYRA by the OeAW, as well as the Boehringer Ingelheim Stiftung. GW acknowledges support from the European Research Council (ERC) (CoG grant #851478); SL acknowledges support from an EMBO Postdoctoral Fellowship (ALTF 236-2024). KT, PD, JH, KD and PH further acknowledge the OP JAK project “Photomachines” Reg. No CZ.02.01.01/00/22_008/0004624 (Czech Ministry of Education, Youth, and Sports (MEYS)).

## Author contribution

F. G. and A.V. designed the study. F.G., S.L., P.V., D.D., Y.T., K.D., L.E., performed experiments. P.H., D.T. and J.H. isolated and purified Nostatin A. P.H. and A.V. secured funding. P.H., M.U., M.A., S.L. and G.W. provided critical feedback for the design and implementation of chemoproteomics experiments. F.G. prepared figures, F.G. and A.V. wrote the manuscript with the input of all co-authors.

---

**Supplementary Figure 1:**
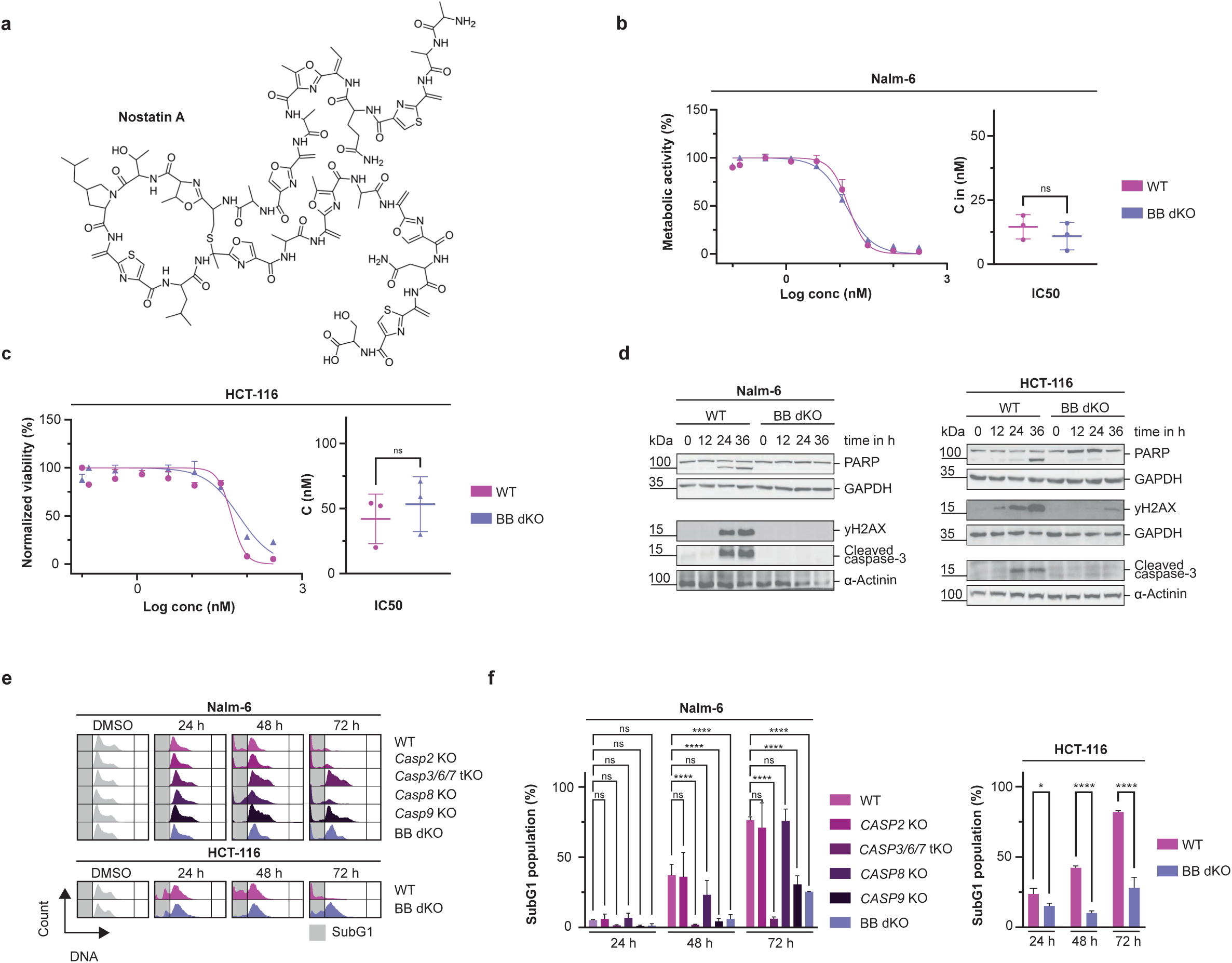
Nostatin A induces mitochondrial apoptosis across different cancer cell lines. a. Chemical structure of Nostatin A b. Metabolic activity of Nalm-6 WT and BAX/BAK dKO cells treated with graded concentrations of NoA for 72 h was assessed using the CellTiter-Glo® Luminescent Cell Viability Assay. Data points represent mean ± SD of three technical replicates. IC₅₀ values are shown as mean ± SD of N = 3 biological replicates and were determined by linear regression. Statistical significance between IC₅₀ values was assessed using an unpaired t-test. ns = not significant. c. Metabolic activity of HCT-116 WT and BAX/BAK dKO cells treated with graded concentrations of NoA for 72 h was assessed using the CellTiter-Glo® Luminescent Cell Viability Assay. Data points represent mean ± SD of three technical replicates. IC₅₀ values are shown as mean ± SD of N = 3 biological replicates and were determined by linear regression. Statistical significance between IC₅₀ values was assessed using an unpaired t-test. ns = not significant. d. Western blot analysis of Nalm-6 and HCT-116 WT and BB dKO cells treated with 50 nM NoA. Cells were harvested at 0, 12, 24, and 36 h after treatment, and protein lysates were probed with the indicated antibodies. e. Representative cell cycle profiles of different Nalm-6 and HCT-116 clones after 24, 48 and 72 h of treatment with 50nM of NoA. f. Quantification of SubG1 populations in Nalm-6 and HCT-116 clones. Bars represent mean ± SD of N = 3 biological replicates. Statistical significance was determined by two-way ANOVA with Bonferroni correction for multiple comparisons, comparing SubG1 percentages of WT and KO clones at each time point. *P < 0.05, ****P < 0.0001.

**Supplementary Figure 2:**
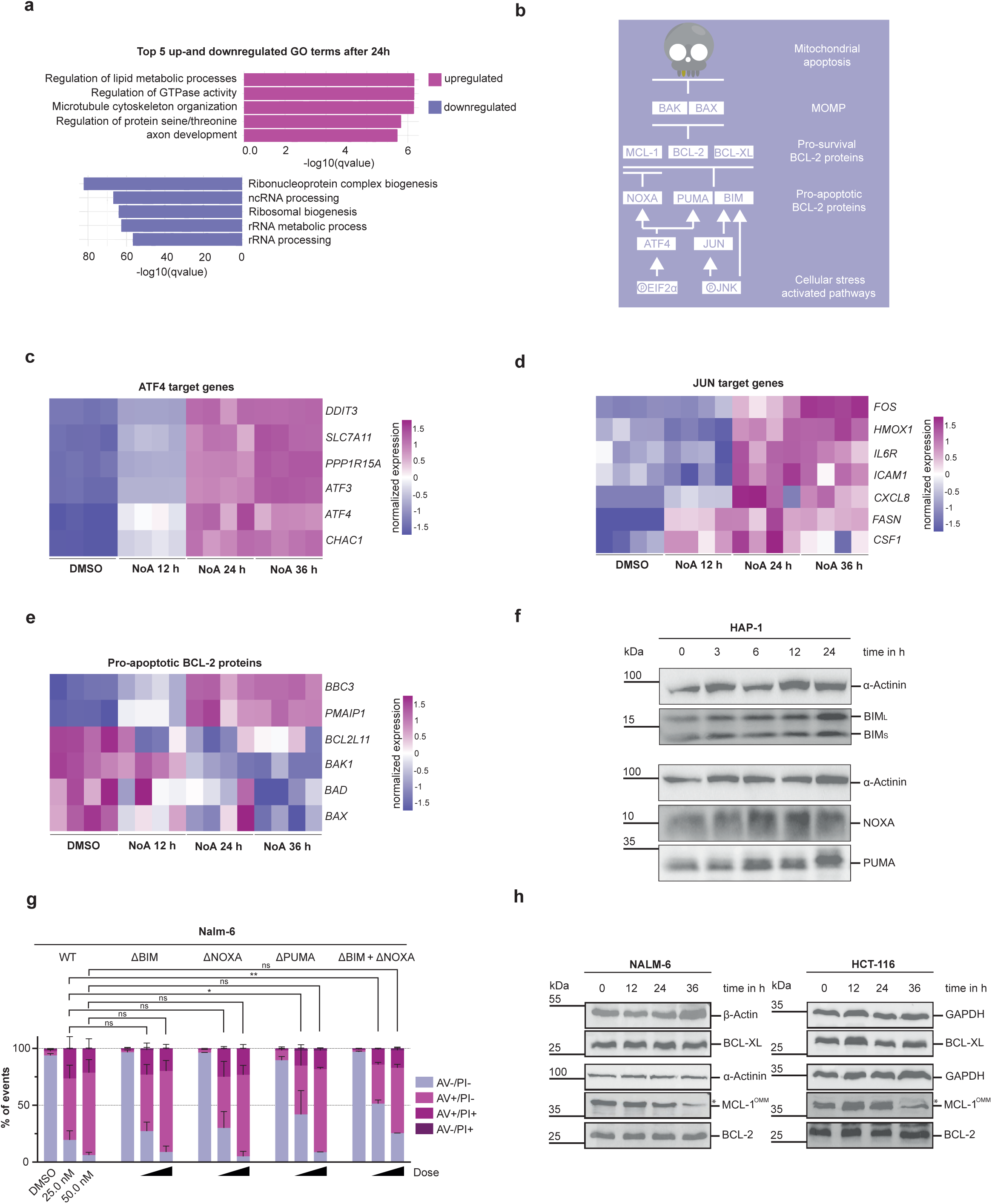
Nostatin A induces ATF4- and JUN-dependent transcriptome changes preceding apoptosis. a. Gene ontology analysis of differentially expressed genes of HAP-1 cells after 24 h treatment with 50 nM of NoA or DMSO vehicle control. b. Schematic of mitochondrial apoptosis regulation by BCL-2 family members. Chronic cellular stress BH3-only proteins upregulation antagonize pro-survival BCL-2 members and activate BAX/BAK, resulting in mitochondrial outer membrane permeabilization and apoptosis. c. Heatmap showing regulation of canonical ATF4 target genes extracted from the RNA-seq data. Values show row scaled vst-normalized expression of differentially expressed genes. Values of n=4 technical replicates are shown. d. Heatmap showing regulation of canonical JUN target genes extracted from the RNA-seq data. Values show row scaled vst-normalized expression of differentially expressed genes. Values of n=4 technical replicates are shown. e. Heatmap showing regulation of pro-apoptotic BH-3 proteins extracted from the RNA-seq data. Values show row scaled vst-normalized expression of differentially expressed genes. Values of n=4 technical replicates are shown. f. Western blot analysis of HAP-1 WT cells treated with 50 nM NoA. Cells were harvested at 0, 3, 6, 12 and 24 h after treatment, and protein lysates were probed with the indicated antibodies. g. Nalm-6 WT and KO clones were treated with 12, 25 and 50 nM of NoA. Annexin V surface binding and Propidium Iodide uptake was measured after 72 h to assess viability of the cells. Bar plots show mean ± SD of three biological replicates. Legend: AV−/PI− = live cells, AV+/PI− = early apoptosis, AV+/PI+ and AV−/PI+ = late apoptosis. Statistical significance was determined using a two-way ANOVA with Bonferroni correction for multiple comparisons, comparing percentages of live cells of corresponding concentrations between WT and KO clones. \**P* < 0.05, \*\**P* < 0.005 h. Western blot analysis of Nalm-6 and HCT-116 WT cells treated with 50 nM NoA. Cells were harvested at 0, 12, 24 and 36 h after NoA treatment and protein lysates probed with indicated antibodies. MCL-1 OMM refers to the full-length, anti-apoptotic isoform MCL-1.

**Supplementary Figure 3:**
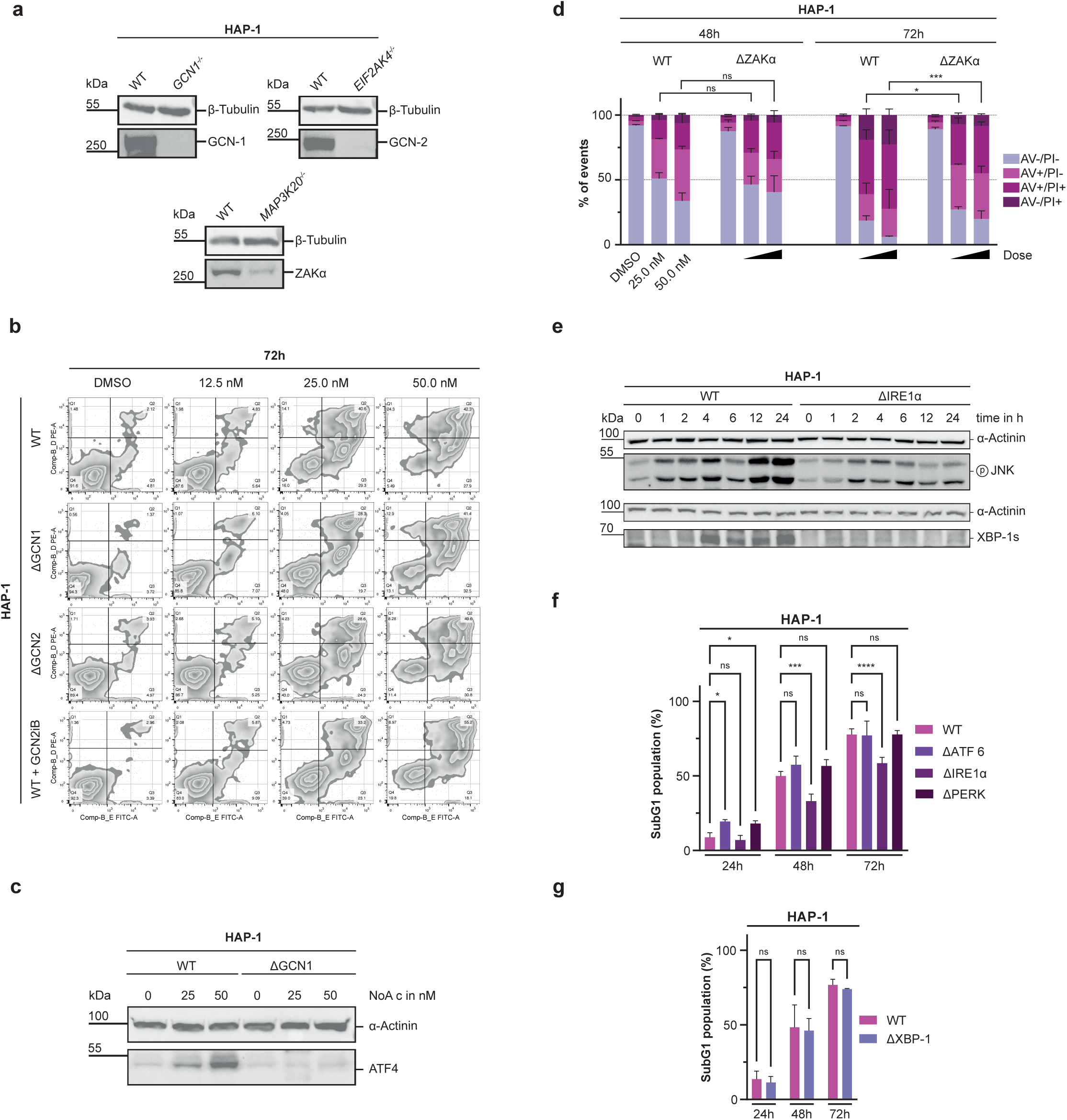
NoA activates the ISR and ribotoxic stress response through GCN1/GCN2 and ZAKα. a. WB analysis to validate HAP-1 KO clones obtained from Horizon Genomics. b. Representative FACS plots of HAP-1 WT, GCN1 KO, and GCN2 KO cells treated with the indicated concentrations of NoA for 72 h. HAP-1 WT cells co-treated with 10 μM GCN2iB for 72 h are also shown. Values indicate percentages of total events within that quadrant. c. HAP-1 WT and ZAKα KO clones were treated with 25 and 50nM of NoA. Annexin V surface binding and Propidium Iodide uptake was measured after 48 and 72h to assess cell viability. Bar plots show mean ± SD of three biological replicates. Legend: AV−/PI− = live cells, AV+/PI− = early apoptosis, AV+/PI+ and AV−/PI+ = late apoptosis. Statistical significance was determined using a two-way ANOVA with Bonferroni correction for multiple comparisons, comparing percentages of live cells of corresponding concentrations between WT and ZAKα KO clones. \**P* < 0.05, \*\*\**P* < 0.001 d. WB analysis comparing HAP-1 WT and IRE1α KO cells after treatment with 50 nM of NoA. Samples were collected at indicated timepoints and lysates were probed with the indicated antibodies. e. WB analysis comparing ISR activation in HAP-1 WT and GCN1 KO cells after 24h treatment with 25 and 50nM of NoA. Lysates were probed with the indicated antibodies. f. Quantification of SubG1 populations in HAP-1 clones treated with 50 nM of NoA for indicated durations. Bars represent mean ± SD of N = 3 biological replicates. Statistical significance was determined by two-way ANOVA with Bonferroni correction for multiple comparisons, comparing SubG1 percentages of WT and KO clones at each time point. *P < 0.05, \*\*\**P* < 0.001, ****P < 0.0001. g. Quantification of SubG1 populations in HAP-1 WT and XBP-1 KO cells treated with 50 nM of NoA for indicated durations. Bars represent mean ± SD of N = 3 biological replicates. Statistical significance was determined by two-way ANOVA with Bonferroni correction for multiple comparisons, comparing SubG1 percentages of WT and KO clones at each time point.

**Supplementary Figure 4:**
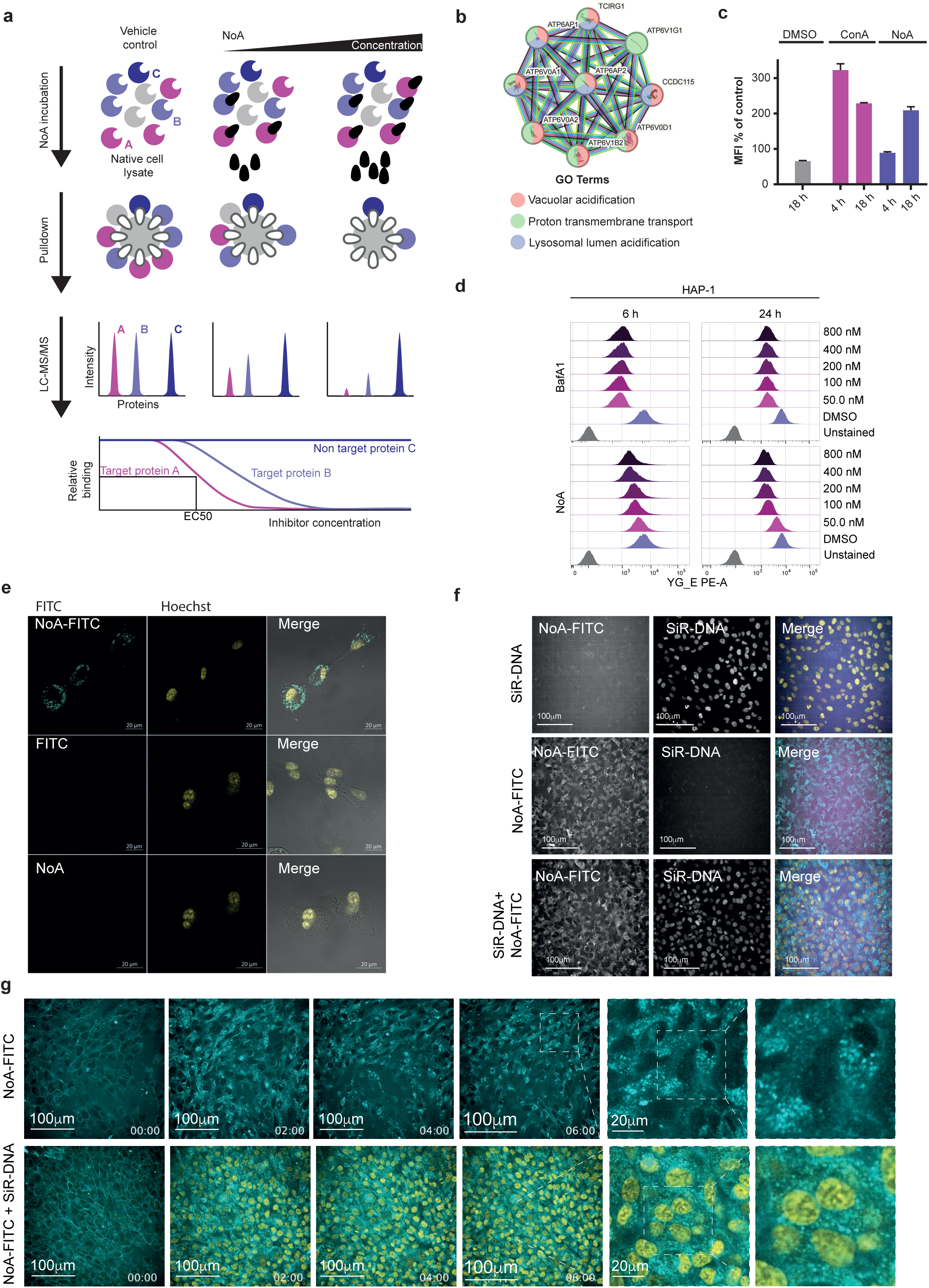
Nostatin A binds and inhibits V-type H⁺-translocating ATPase. a. Schematic representation of the chemoproteomic competition binding assay. b. Protein-protein interaction network of significantly depleted targets identified as NoA interactos. Individual proteins are represented as nodes, while predicted or established interactions between them are represented as edges. Network and GO term analysis were generated using the STRING database (version 12.0). c. Assessment of lysosomal acidification in cells treated with NoA or the V-ATPase inhibitor concanamycin A in MDA-MB 231 cells. Cells were treated with 100 nM NoA or 50 nM concanamycin A for 4 h or 18 h, and lysosomal pH was measured using a pH-sensitive fluorescent dextran probe. Fluorescence intensity increases upon lysosomal deacidification and was quantified by flow cytometry as mean fluorescence intensity (MFI). Bar plots represent values of two biological replicates. d. Representative flow cytometry histograms of LysoTracke fluorescence in HAP-1 WT cells treated with NoA or bafilomycin A1 (BafA1) at the indicated concentrations for 6 or 24 h. Fluorescence intensity reflects lysosomal acidification. DMSO-treated cells served as the control. e. Confocal fluorescence microscopy images of RPE-1 cells treated with 100 nM NoA-FITC for 6 h. NoA-FITC is shown in cyan and nuclei stained with Hoechst in yellow. Images are shown with or without the nuclear stain. f. Confocal fluorescence microscopy images of RPE-1 cells treated with 100 nM NoA-FITC for 24 h. NoA-FITC is shown in cyan and nuclei stained with SiR-DNA in yellow. Images are shown with or without the nuclear stain. g. Confocal fluorescence microscopy time-course imaging of RPE-1 cells treated with 100 nM NoA-FITC. Cells were imaged after 2, 4, and 6 h of treatment. NoA-FITC is shown in cyan and nuclei stained with SiR-DNA in yellow. Images are shown for cells treated with NoA-FITC alone or co-stained with SiR-DNA.

**Supplementary Figure 5:**
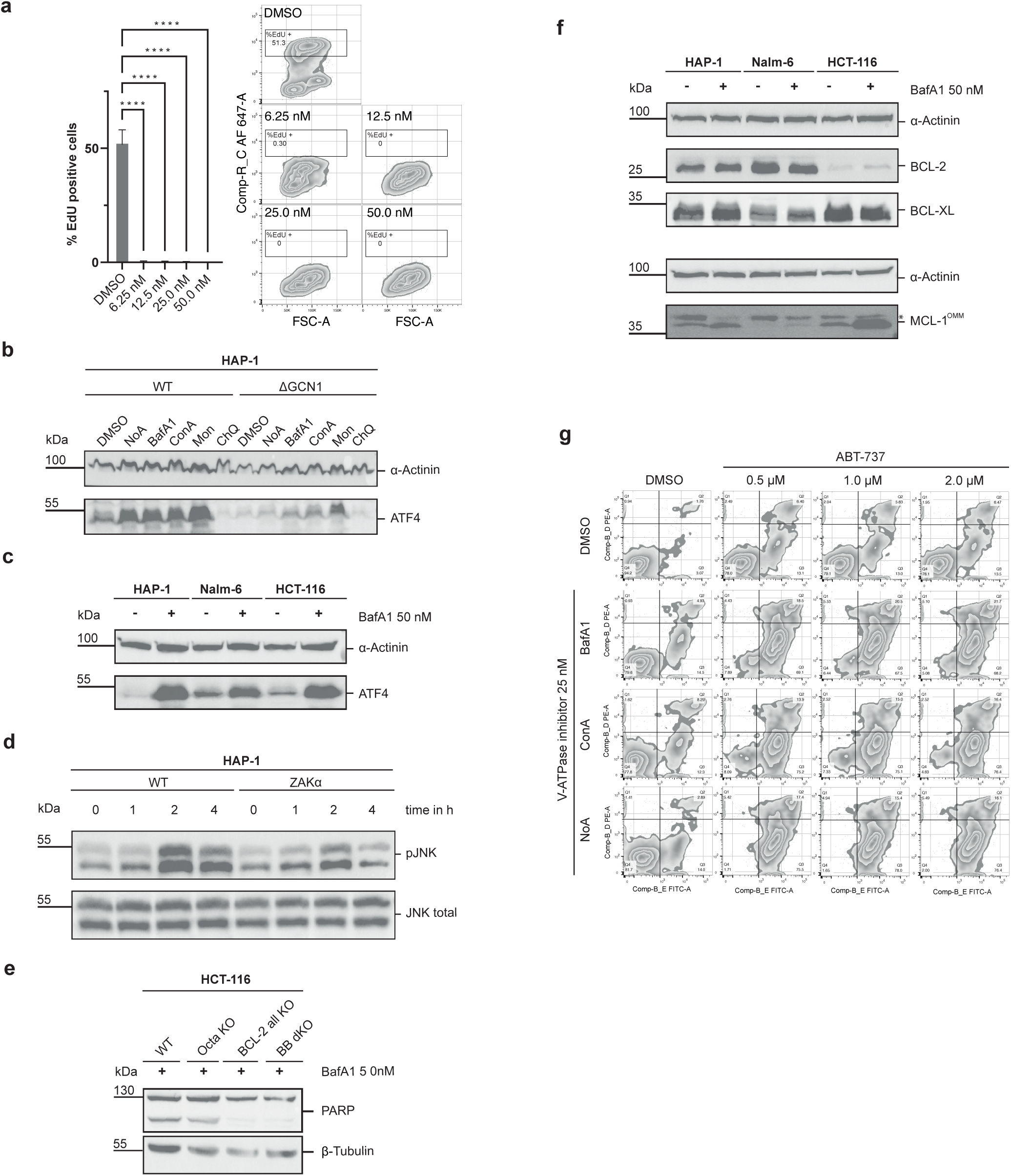
V-ATPase inhibition activates the RSR/ISR and synergizes with BH3 mimetic treatment. **a.** EdU incorporation in HAP-1 WT celle after 24 h treatment with graded concentrations of BafA1. Bar graphs (left) are showing means ± SD of N=3 biological replicates of % of EdU positive cells with representative dot plots (right). Statistical significance was assessed using a one-way ANOVA with Dunnet’s multiple comparisons test. \*\*\*\**P* < 0.0001 **b.** WB analysis of HAP-1 WT, GCN1 KO cells treated with NoA, BafA1, ConA (50 nM) Monensin sodium (2 μM) or Chloroquine (10 μM) for 24 h. Cell lysates were probed for the indicated proteins. **c.** WB analysis of HAP-1, Nalm-6 and HCT-116 WT cells treated with 50 nM of BafA1 for 24h. Cell lysates were probed for the indicated proteins. **d.** WB analysis comparing HAP-1 WT and ZAKα KO cells after treatment with 50 nM of BafA1. Samples were collected at indicated timepoints and lysates were probed with the indicated antibodies. **e.** WB analysis of HCT-116 WT, Octa KO, BCL-2 all KO and BB dKO cells after 24 h of treatment with 50nM of BafA1. **f.** WB analysis of HAP-1, Nalm-6 and HCT-116 WT cells treated with 50nM of BafA1 for 24 h. Cell lysates were probed for the indicated proteins. MCL-1 OMM refers to the full-length, anti-apoptotic isoform MCL-1. **g.** Representative FACS plots of HAP-1 cells treated with indicated V-ATPase inhibitors (25 nM) alone or in combination with ABT737 (0.5, 1 and 2 μM) for 24 h. Values indicate percentages of total events within that quadrant.

